# Comparative proteomics of osmotic signal transduction mutants in *Botrytis cinerea* explain loss of pathogenicity phenotypes and highlight interaction with cAMP and Ca^2+^ signalling pathways

**DOI:** 10.1101/557983

**Authors:** Jaafar Kilani, Marlène Davanture, Michel Zivy, Sabine Fillinger

## Abstract

- Signal transduction (ST) is essential for rapid adaptive responses to changing environmental conditions through rapid post-translational modifications of signalling proteins and downstream effectors that regulate the activity of target proteins and/or the expression of downstream genes.
- We have performed a comparative proteomics study of ST mutants in the phytopathogenic fungus *Botrytis cinerea* during axenic growth under non-stressed conditions to decipher the roles of two kinases of the hyper-osmolarity pathway in *B. cinerea* physiology. We studied the mutants of the sensor histidine kinase Bos1 and of the MAP kinase Sak1.
- Multiplex shotgun proteomics detected 628 differential proteins between mutants and wild-type, 280 common to both mutants, indicating independent and shared regulatory functions for both kinases. Gene ontology analysis showed significant changes in proteins related to plant infection (secondary metabolism enzymes, lytic enzymes, proteins linked to osmotic, oxidative and cell wall stress) that may explain the virulence defects of both mutants. Intracellular accumulation of secreted proteins in the *Δbos1* histidine-kinase mutant suggests a potential secretion defect. The proteome data also highlight a new link between Sak1 MAPK, cAMP and Ca^2^+ signalling.
- This study reveals the potential of proteomic analyses of signal transduction mutants to decipher their biological functions.

## INTRODUCTION

*Botrytis cinerea* is a necrotrophic, polyphageous plant pathogenic fungus responsible for grey mold disease of nearly 600 plant genera (Elad *et al.*, 2016), many of which have significant agronomic importance. *B. cinerea* can infect different plant organs such as leaves, flowers or fruits and causes extensive pre- and post-harvest damage leading to yield losses. The resulting economic impact may then be considerable (Droby & Lichter, 2007; Romanazzi *et al.*, 2016). The infection process of *B. cinerea* involves the secretion of lytic enzymes, production of toxins, reactive oxygen species and small RNAs (González *et al.*, 2016). In return, *B. cinerea* has to face to many stresses caused by its environment and in particular by the infected plant such as osmotic, oxidative and cell wall stress, phytoalexins and other plant defence compounds (Windram *et al.*, 2016). Response and adaptation to these stresses is therefore crucial for the survival of the fungus. To be able to persist in this hostile environment and to counter the plant defence, the fungus develops rapid adaptive responses. It perceives these stresses and transmits them to the cell in order to give an adequate response *via* signalling pathways (Schumacher, 2016b).

One of the main signalling pathways in eukaryotes involves MAPKs (Mitogen Activated Protein Kinase), evolutionary conserved signalling pathways. Depending on their phosphorylation status, the activated MAPKs act by regulating numerous cellular processes, notably by activating transcription factors (Yang *et al.*, 2003). MAPK cascades are preserved in ascomycetes, but their roles differ according to the fungal species (Hamel *et al.*, 2012). In *Saccharomyces cerevisiae,* five MAPK pathways have been described controlled respectively by Fus3, Kss1, Slt2, Hog1 and Smk1 (Gustin *et al.*, 1998). The functionally redundant MAPKs Fus3 and Kss1 are involved in sexual and asexual reproduction. Slt2 regulates cell cycle progression and cell wall integrity, while Hog1 regulates the response to osmotic and oxidative stresses. The less studied MAPK Smk1 is involved in the assembly of the ascospore wall. Some MAPK pathways have partially overlapping functions or are interconnected (Fuchs & Mylonakis, 2009). With the exception of Smk1, homologues of these MAPKs are found in filamentous fungi (Hamel *et al.*, 2012). Thus, the *B. cinerea* MAPKs Bmp1, Bmp3 and Sak1 are respectively the homologues of the yeast proteins Fus3/Kss1, Slt2 and Hog1 (reviewed in (Schumacher, 2016b)).

In *B. cinerea,* the MAPK Bmp1 is involved in vegetative growth, hydrophobic surface perception, pathogenicity, spore and sclerotium formation (Zheng *et al.*, 2000; Doehlemann *et al.*, 2006; Schamber *et al.*, 2010). The MAPK Bmp3 is involved in adaptation to hypo-osmotic and oxidative stress and to the fungicide fludioxonil (Rui & Hahn, 2007).

The Sak1 MAPK pathway controls fungal development such as growth and macroconidia formation. It is involved in the adaptation to various stresses such as osmotic (ionic), oxidative and cell wall stress but also in infection (Segmuller *et al.*, 2007; Liu *et al.*, 2011). Indeed, the *Δsak1* mutant is unable to form infection structures and thus to penetrate its host unless the latter is injured (Segmuller *et al.*, 2007). Cross-talk between the Bmp3 and Sak1 pathways has been detected during oxidative stress (Liu *et al.*, 2011).

Sak1 targets transcription factors such as BcReg1 and BcAtf1 (Michielse *et al.*, 2011; Temme *et al.*, 2012). BcReg1 is involved in the regulation of secondary metabolite synthesis, sporulation and host tissue colonization (Michielse *et al.*, 2011) while BcAtf1 is involved in sporulation and the regulation of genes involved in stress response (Temme *et al.*, 2012). Upstream of the osmoregulatory MAPK pathway, the perception and transmission of signals pass through a network of signaling proteins referred to as a two-component systems. In fungi, the histidine kinase sensor is a hybrid protein (HHK) that contains both a sensor domain and a response regulatory (RR) domain (Herivaux *et al.*, 2016). Thus, following the perception of a signal, the histidine kinase catalyses its autophosphorylation on the conserved histidine residue, then transfer of the phosphate group to a conserved aspartate residue of the RR domain. The signal is then transmitted via a phosphotransfer protein (PTH) to a RR protein (Herivaux *et al.*, 2016). In *B. cinerea,* there are two histidine kinases upstream of the Sak1 pathway, Bos1 and BcHK5. The class VI histidine kinase, BcHK5, is the homologue of the unique HHK in *S. cerevisiae* Sln1. Indispensable in yeast because of its role in regulating stress response, this histidine kinase is not essential for development and pathogenicity in *B. cinerea* (reviewed in (Schumacher, 2016b)). In several fungi, the deletion of class III histidine kinase leads to resistance to phenylpyrroles and dicarboximides but also to sensitivity to osmotic, oxidative and parietal stresses (Ochiai *et al.*, 2001; Avenot *et al.*, 2005; Motoyama *et al.*, 2005). This holds true also in *B. cinerea* (Viaud *et al.*, 2006). An original finding was the the constitutive phosphorylation of the MAPK Sak1 in the *Δbos1* mutant, indicating that the two-component system negatively controls the activation of the osmoregulatory pathway (Liu *et al.*, 2008).

Heller et al (Heller *et al.*, 2012) conducted a small scale transcriptomic analysis of the *B. cinerea Δsak1* mutant compared to the wild type. Albeit performed on a limited set of genes, this macroarray analysis revealed Sak1 as central element in the overall response to stress and in the infectious process at the transcriptional level.

Another level in systematic analysis is achieved through proteomics (Breker & Schuldiner, 2014). The analysis of the accumulation of intracellular proteins takes into account all regulations during protein synthesis (i.e., transcription and translation) and degradation (mRNA and protein turn-over).

In this study, we analysed the role of the osmotic ST pathway in the biology of *B. cinerea* and particularly the role of the histidine kinase Bos1 and the MAPK Sak1. In order to highlight proteins whose abundance depends on the osmoregulatory pathway, we performed a comparative proteomic analysis of the *Δsak1* and *Δbos1* mutants compared to the wild type strain.

Protein abundance variations corroborated the implication of both protein kinases in the infection process. Moreover, this study highlighted cross-talk between the Sak1 and the cAMP pathway which, on its turn, is involved in pathogenicity as well. Finally, our study revealed different sets of regulation patterns among the detected proteins.

## MATERIALS AND METHODS

### Strains, medium and culture conditions

*B. cinerea* wild-type strain, B05.10 (Büttner *et al.*, 1994), as well as the mutants *Δsak1* and *Δbos1* (Segmuller *et al.*, 2007; Liu *et al.*, 2008) were cultivated on solid Sisler medium (KH2PO4 2 g l^-1^, K2HPO4 1.5 g l^-1^, (NH4)2SO4 1 g l^-1^, MgSO4 7H2O 0.5 g l^-1^, glucose 10 g l^-1^, yeast extract 2 g l^-1^, agar 15 g l^-1^).

Pre-cultures were made from calibrated explants (5 mm) deposited on plate with solid Sisler medium previously covered with a sterile cellophane membrane (Bio-Rad, USA). The cultures were maintained 4 days at 20°C, in darkness. The mycelium was recovered and crushed for 1 minute in 100 mL of Sisler liquid medium using an Ultra Turax. 500 μl of this shred was spread on solid Sisler cellophane coated medium. The cultures were incubated in the dark at 20°C for 2 days. After these 2 days, the samples were collected and frozen in liquid nitrogen and freeze-dried. Four replicates were made for each strain.

### Protein extraction and tryptic digestion

Proteins of the various genotypes were extracted from lyophilized mycelia of exponential growth phase after grinding mycelium in liquid nitrogen. Proteins were precipitated using a precipitation solution (10% trichloroacetic acid (TCA) and 0.07% β-mercaptoethanol, in acetone). After vortex and centrifugation at 9838 g at −20°C, the pellet was rinsed three times in acetone with 0.07% β-mercaptoethanol to remove TCA. After drying, the pellet was weighed and proteins were solubilized at 20 μl per mg in ZUT buffer (Urea 6 M, Thiourea 2 M, DTT 10 mM, Tris-HCl 30 mM pH 8.8, 0.1% of ZALS (ProgentaTM Zwitterionic Acid Labile Surfactants – Protea Bioscience). After 5 min of centrifugation at 14000 g and 25°C, the supernatant was collected.

Protein concentration was measured with the “2D Quant kit” (GE Healthcare Life Sciences), according to the supplier’s instructions. 40 μg of extracted proteins were sampled and the volume adjusted with ZUT buffer to adjust the concentration to 4 μg μl^-1^. Proteins were alkylated with iodoacetamide (final concentration 50 mM) for 45 min at room temperature in the dark. Samples were then diluted with 50 mM ammonium bicarbonate solution to reduce the urea concentration to 1 M. Proteins were digested overnight at 37°C with 800 ng trypsin (Promega). Digestion was stopped by adding 5 μl TFA (Trifluoroacetic acid) at 20%, then incubated 30 min at room temperature to cleave ZALS. The peptide extracts were desalted on C18 column (Strata™XL 100 μm Polymeric Reversed Phase – Phenomenex).

### LC-MS/MS analysis

For each sample 4 μl of the peptide mixture obtained by trypsic digestion were injected and separated by HPLC (High Pressure Liquid Chromatography) Nano LC-Ultra system (Eksigent, United Kingdom). Chromatography uses a 30 cm C18 column (Nanoseparations, Netherlands) and a 99.9% CH3CN + 0.1% formic acid buffer gradient, according to the following steps: 5% to 35% buffer for 110 min; 35% to 95% buffer for 3 min; 95% buffer for 10 min. The injection into the QexactivePLUS mass spectrometer (Thermo Fisher Scientific, USA) was performed from a nano-electrospray source (New Objective, USA).

The whole system was controlled by Xcalibur 4.0 with the following acquisition method: the acquisition of MS spectra was performed over a range m/z = 400 – 1400 with a resolution of 70000. The eight most intense ions (Top 8) underwent HCD (Higher energy collisional dissociation) fragmentation with collision energy of 27%, thus obtaining an MS/MS spectrum acquired at a resolution of 17500. A dynamic exclusion of 40 sec was used.

Peptide identification was performed using X!Tandem software (Craig & Beavis, 2004) (PILEDRIVER 01/04/2015 version) where mass changes were indicated. The mass change corresponding to carbamidomethylation of cysteines (57.02146 Da) was reported as a systematic change. Potential modifications have been added such as oxidation of methionines (15.99491 Da) and acetylation to N-terminal (+42.01056 Da). The mass tolerance of the precursor was 10 ppm while that of the fragment was 0.02 Da. Only one missed cleavage was allowed.

The *B. cinerea* database provided to the software was retrieved from Ensembl (http://fungi.ensembl.org/Botrytis_cinerea/) and previously functionally re-annotated on UseGalaxy from published data (Amselem *et al.*, 2011) and automatic annotations obtained with Pfam (Finn *et al.*, 2010). A database of standard contaminants has also been added. The identified proteins were filtered and grouped with X!Tandem Pipeline v3.4.0 (Langella *et al.*, 2017). Data with a peptide E-value < 0.01, at least two peptides per protein and a protein E-value of 10^-4^ were retained. The FDR (False Discovery Rate) was estimated at 0.02% for peptides and 0% for proteins.

Identified proteins, peptides and their corresponding spectra were deposited in PROTICdb database (Langella *et al.*, 2013) at the following URL http://moulon.inra.fr/protic/botrytis_signalling using the following login: “review” and password (required during the review process only): “review”. Data will soon be accessible under doi 10.15454/1.5506737620833718E12.

### Quantification, statistical analysis and annotation

Quantification was based on the analysis of the MS1 XIC (eXtracted Ion Current) of each of the identified peptides, by using the MassChroQ software v2.2 (Valot *et al.*, 2011). XIC data (peak intensity integration) was used to quantify proteins for which at least two specific peptides were quantified in at least 95% of the samples. For proteins that did not reach this criterion, detected peaks were counted (peak counting, PC) instead of being integrated.

For XIC quantification, normalization was performed according to the median sample/reference ratio of peptide ion XIC values, the reference being a WT sample. Peptide ion missing XIC data were imputed according to the correlation between peptides of the same protein. The relative abundance of each protein was computed by summing the intensities of its specific peptides. An analysis of variance (linear model) on log_10_-transformed abundances was performed to study the variations according to the strain. Proteins with adjusted p-value < 0.01 and a fold-change (mutant/WT) above 1.5 or below 0.66 were considered significantly affected by the mutant.

For PC, an analysis of variance was performed by using a general linear model and the same criteria as those employed for XIC quantification were used to consider a protein significantly affected by the mutation. The R software (version 3.4.4 (R Core Team, 2018)) was used for all statistical analyses.

After grouping proteins into regulatory groups, functional enrichment according to biological processes was analysed with Blast2GO (Gotz *et al.*, 2008) for each group. This enrichment uses an exact Fisher test with a p-value < 0.05. This same type of test was also used to perform manual enrichment studies from a list of selected biological functions.

### Intracellular cAMP measurements

The different strains were grown under conditions identical to those for the proteomic analyses. Intracellular cAMP concentration was measured using the Amersham cAMP Biotrak Enzyme-immunoassay System kit (GE Healthcare). 4 mg of lyophilized mycelium was crushed in a Fast-Prep machine (MP Biomedicals) using Garnet Matrix A (MP Biomedicals) and Lysis Buffer 1B (supplied in the assay kit). After centrifugation, the supernatant was collected and used for the enzyme test. The calculation of the cAMP concentration was performed against a standard curve plotted using a four-parameter logistic regression method. The cAMP concentrations were normalized to the nucleic acid concentration in the extract, measured at λ=260 nm and expressed as fmoles ng^-1^ of nucleic acids.

## RESULTS

### Quantitative proteomic analysis of signal transduction mutants

*Δbos1* and *Δsak1* mutants’ protein content were compared to the parental wild-type strain B05.10 exponentially grown under axenic *in vitro* conditions. After tryptic digestion and peptide analysis by LC-MS/MS, protein identification and quantification were performed. 67% of the detected spectra could be assigned to peptides. 2425 proteins out of the 11701 predicted *B. cinerea* proteins (20.7% of the theoretical proteome) were detected in the total dataset. XIC analysis allowed to obtain quantitative profiles of 1263 proteins out of the previously mentioned 2425 (52%) and the others were analysed by peak counting (PC).

### Identification of differentially abundant proteins between wild-type and osmosensing mutants

In the *Δbos1* mutant, the abundance of 417 proteins (377 in XIC and 40 in PC) differed significantly from the wild-type, while in the *Δsak1* mutant the abundance of 481 proteins (419 in XIC and 62 in PC) differed significantly from the wild-type. 270 proteins were common to both mutants (Figure 1).

**Figure 1:**
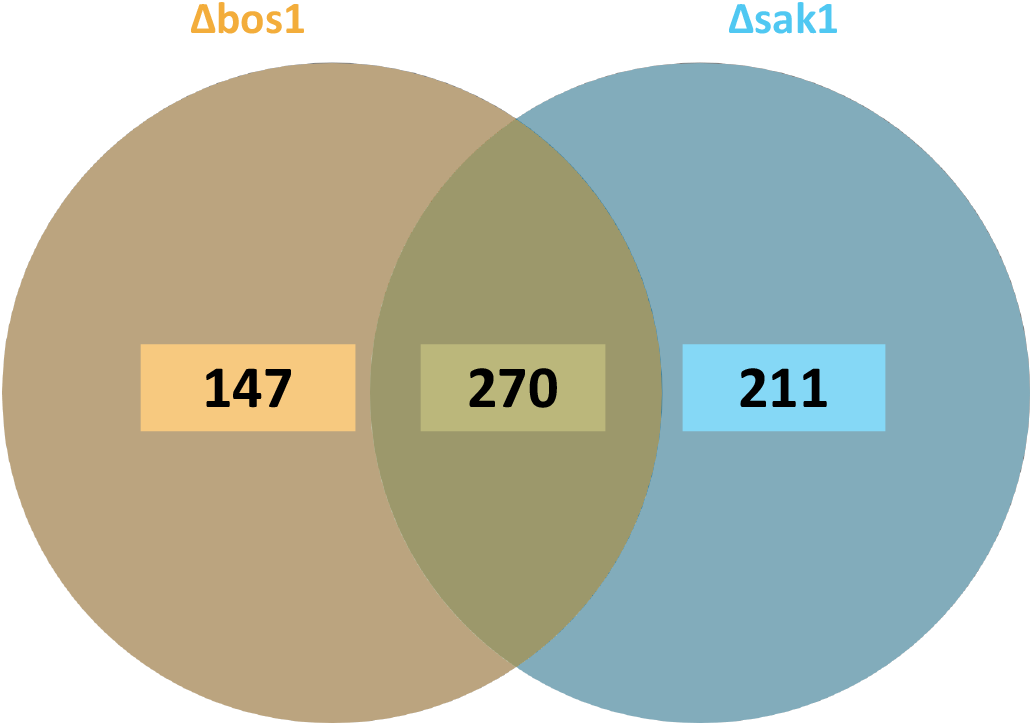
Differentially accumulating proteins in *Botrytis cinerea Δbos1* and *Δsak1* mutants. The number indicated proteins whose abundance is significantly modified in *Δbos1* and *Δsak1* compared to the wild-type, in steady state conditions.

Table 1 recapitulates the number of proteins whose abundance varied significantly in both mutants jointly or independently. Among the 243 proteins similarly affected in both mutants, 143 were found less abundant and 100 more abundant than in the wild-type, reflecting positive or negative regulation respectively. The number of proteins whose abundance variations were contrasting in both mutants, and consequently following potentially opposing regulation patterns, was considerably lower (n=27). We further detected proteins controlled independently by either Sak1 (88 positively and 123 negatively) or Bos1 (89 positively, 58 negatively).

**Table 1:** Functional categories of differentially regulated proteins according to dependence on Bos1 and Sak1 respectively. “+” indicates positive regulation, “-” indicates negative regulation.

| Type of regulation | Blast2GO enrichment | Enrichment (manually annotated) | Number of identified proteins |
| --- | --- | --- | --- |
| Bos1(+)/Sak1(+) | Nucleic acid metabolism<br>Carbohydrate metabolic process<br>Pyruvate metabolic process<br>Lactate metabolic process<br>Gluconeogenesis | Carbohydrate-Active enZymes (CAZymes)<br>Secreted proteins<br>Secondary metabolism | 143 |
| Bos1(-)/Sak1(-) | Alpha-amino acids biosynthetic process | Secondary metabolism<br>Oxidative stress response<br>Respiration alternative | 100 |
| Bos1(+)/Sak1(-) | *nd* | Secondary metabolism | 13 |
| Bos1(-)/Sak1(+) | *nd* | Secreted proteins<br>Carbohydrate-Active enZymes | 14 |
| Bos1(+) | Glycoprotein metabolic process<br>Macromolecule glycosylation | Carbohydrate-Active enZymes<br>Primary metabolism | 89 |
| Bos1(-) | Nucleotide metabolic process<br>Purine nucleobase metabolic process<br>Cellular amino acid metabolic process | Proteases | 58 |
| Sak1(+) | Monosaccharide metabolic process<br>Oxoacid metabolic process<br>Cell redox homeostasis | Oxidative stress response<br>Secreted proteins<br>Detoxification of ROS | 88 |
| Sak1(-) | Carboxylic acid metabolic process<br>Coenzyme biosynthetic process | Secondary metabolism<br>Proteases<br>Signalling pathway components | 123 |

### Enrichment study of functional groups

A functional enrichment analysis based on GO terms and manual annotation was conducted to identify biological processes enriched in each group. Bos1 and Sak1 play a positive role on nucleic acid metabolism, energy metabolism, CAZYmes (Carbohydrate-Active enZYmes), secreted proteins, and secondary metabolism. On the opposite, they negatively regulate mainly the biosynthesis of alpha-amino acids, the response to oxidative stress, alternative respiration and distinct pathways of secondary metabolism. In the groups inversely regulated by Bos1 and Sak1, manual enrichment analysis revealed proteins involved in secondary metabolism, as well as secreted proteins and CAZYmes. The group regulated positively only by Bos1 is enriched in proteins involved in glycosylation of macromolecules, primary metabolism, as well as some CAZYmes. In a negative way, Bos1 mainly regulates proteins of nucleotide and amino acid metabolism.

The group of proteins positively regulated by Sak1 alone is enriched in secreted proteins, as well as proteins involved in the response to oxidative stress but also in the production of ROS. Proteins negatively regulated by Sak1 alone comprise a majority of proteases, as well as proteins involved in secondary metabolism and signalling pathways.

In conclusion, the majority of regulatory groups are enriched in proteins of various functions, all of which participate to the infection process in *B. cinerea*. The functional groups linked to infection are detailed in the following paragraphs.

### Regulation of secondary metabolism

Secondary metabolites are important players during the infection of plants by *B. cinerea* (Collado & Viaud, 2016). Astonishingly, our proteomic analysis revealed contrasting regulatory classes for secondary metabolism enzymes.

Analysis of the Bot proteins (Table 2) involved in botrydial synthesis (Rossi *et al.*, 2011) revealed an increase in the abundance of BcBot4 (cytochrome P450 monooxygenase), BcBot2 (sesquiterpen cyclase), and BcBot3 (cytochrome P450 monooxygenase) in the *Δsak1* mutant, while these proteins were less abundant in the *Δbos1* mutant. The abundance of BcBot1 (cytochrome P450 monooxygenase) did not vary significantly in the *Δsak1* mutant but decreased in the *Δbos1* mutant. The abundance of proteins of the *Bcbot* gene cluster is positively regulated by the histidine kinase Bos1 while MAPK Sak1 negatively regulates several of them, indicating the regulatory roles of both proteins in botrydial biosynthesis. Analysing at the *Bcboa* cluster (Table 2) involved in botcinic acid toxin synthesis (Dalmais *et al.*, 2011), only BcBoa1 (putative NmrA-like regulator) and BcBoa17 (putative dehydrogenase) were detected as significantly different compared to the wild-type.” BcBoa1 was more abundant in the *Δbos1* mutant compared to the wild-type, while its accumulation was reduced in the *Δsak1* mutant. On the opposite, BcBoa17 was found more abundant only in the *Δsak1* mutant than in the wild-type.

**Table 2:**
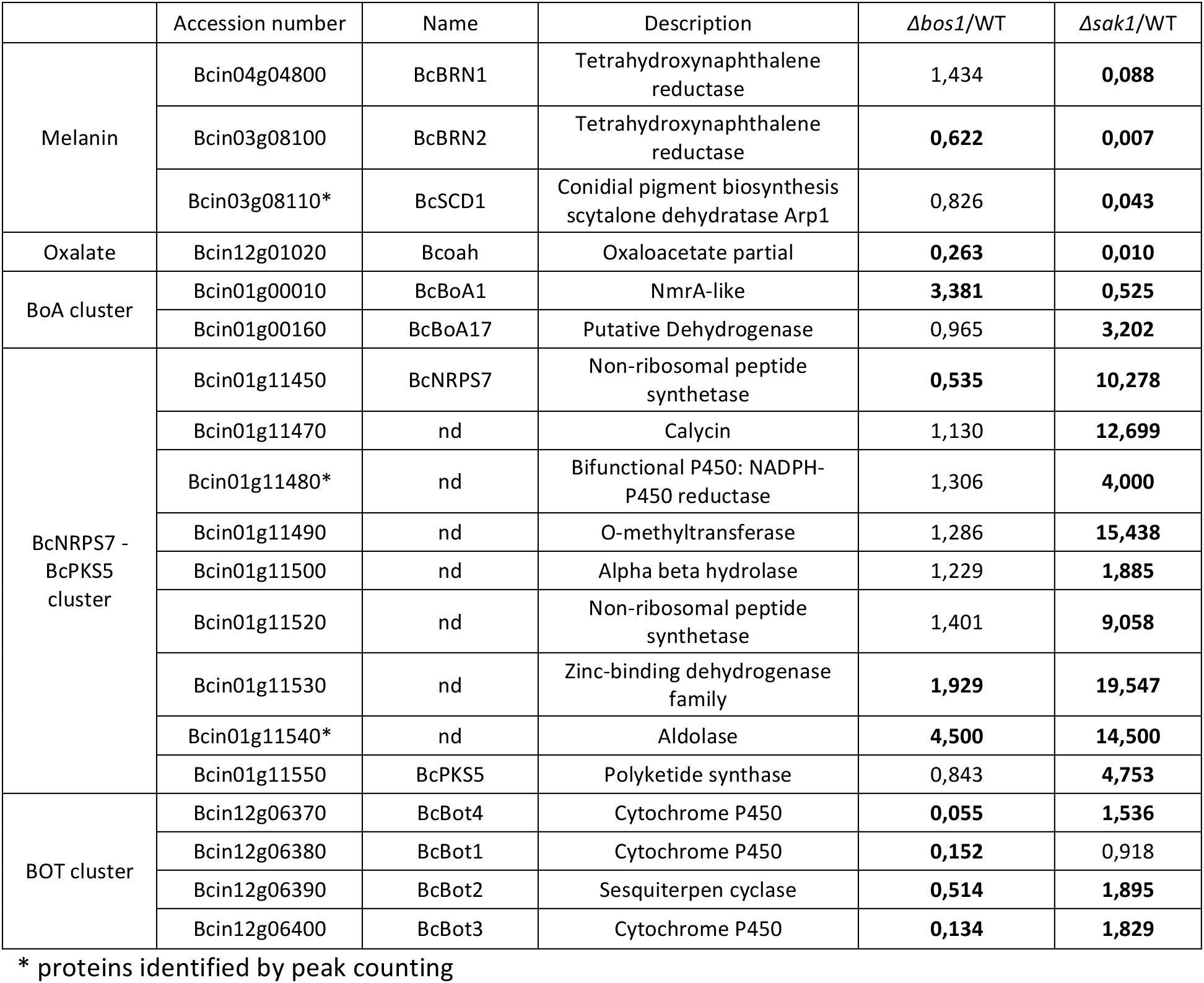
Differentially produced secondary metabolism proteins in *Δbos1* and *Δsak1 mutants.* Bold numbers indicate over threshold values.

Among the other secondary metabolism proteins (Table 2), nine proteins of the BcNRPS7-BcPKS5 cluster (Schumacher *et al.*, 2015) were detected in our proteomic analysis: the non-ribosomal peptide synthetase BcNRPS7, calycin, O-methyltransferase, alpha/beta hydrolase, methyltransferase, enoyl-reductase, aldolase and the polyketide synthase BcPKS5. All nine proteins were identified as being more abundant in the *Δsak1* mutant compared to the wild-type. In the *Δbos1* mutant the abundance of three of these proteins also varied, that of enoyl-reductase and aldolase was found increased but to a lesser extent than in the *Δsak1* mutant. These results highlighted negative regulation of the BcNRPS7-BcPKS5 cluster by MAPK Sak1 and mostly independent of Bos1.

Proteins involved in oxalic acid or melanin biosynthesis (Table 2) were found affected in the osmosensensing mutants. Especially in *Δsak1,* the abundance of oxaloacetate acetylhydrolase decreased considerably, but the abundance of three key enzymes of melanin biosynthesis was highly impacted as well. Indeed, the tetrahydroxynaphtalene reductases BcBrn1 and BcBrn2, the Scytalone dehydratase BcScd1 (Schumacher, 2016a) were less abundant in the MAPK mutant while only BcBrn2 abundance was reduced in the *Δbos1* mutant.

### Proteins involved in osmotic, oxidative and cell wall stress response

#### Osmotic stress

Considering the proteome of osmosensing signal transduction mutants, we were particularly interested in functions related to osmotic stress (Table 3). Glycerol is involved in the osmotic stress response (Blomberg & Adler, 1989) while mannitol is involved in both osmotic and oxidative stress response (Dulermo *et al.*, 2010; Meena *et al.*, 2015). Previous studies showed the requirement for Sak1 in glycerol biosynthesis (Liu *et al.*, 2008). Analysis of our proteomic data with respect to polyol metabolism revealed a reduction in glycerol dehydrogenase and glycerol-3-phosphate dehydrogenase content in both mutants. Regarding mannitol metabolism, we observed a significant decrease in mannitol dehydrogenase and mannitol-1-phosphate 5-dehydrogenase in the *Δsak1* mutant.

**Table 3:**
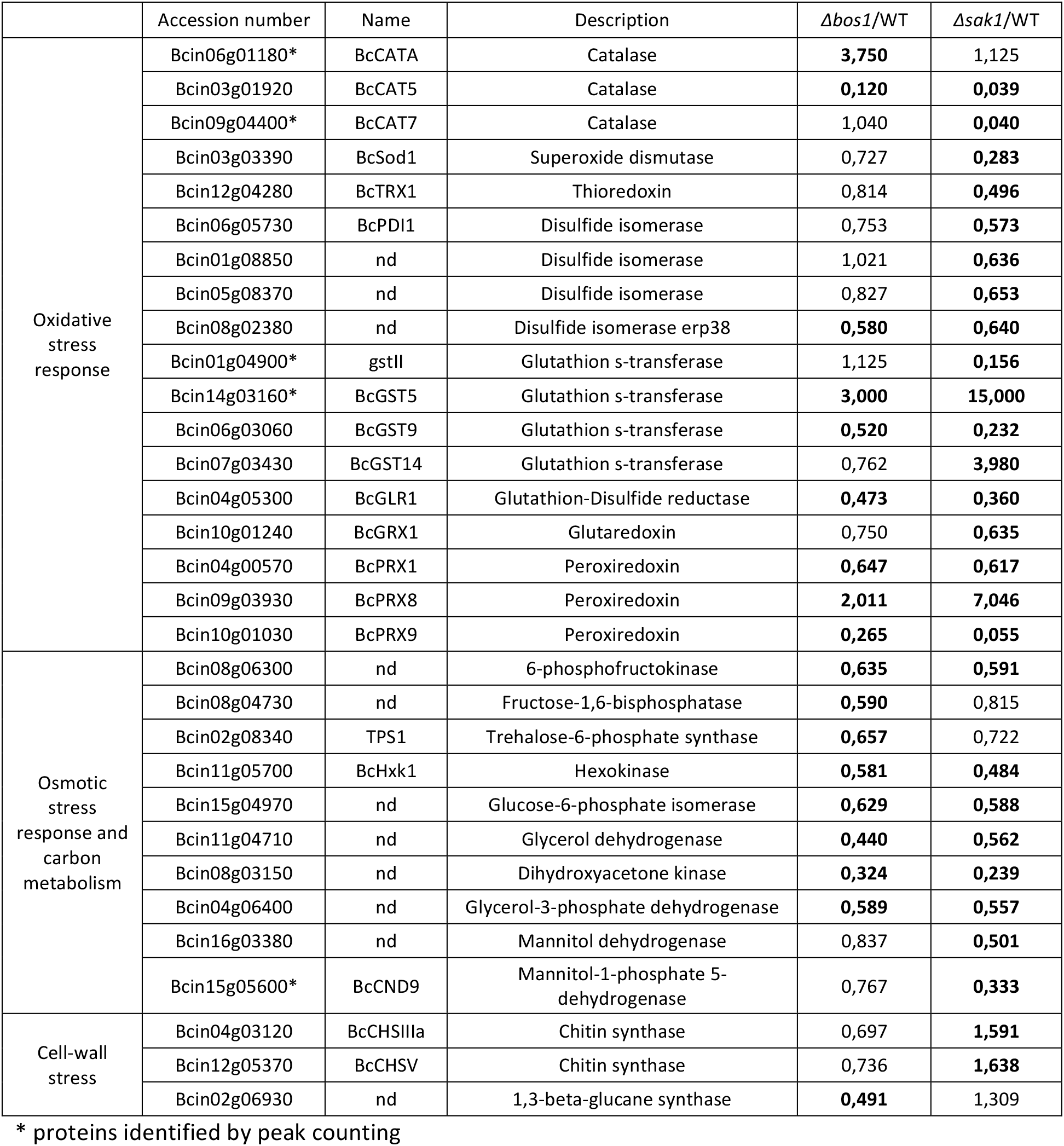
Differentially produced proteins involved in the response to different stresses in *Δbos1* and *Δsak1 mutants* compared to wild-type. Bold numbers indicate over threshold values.

#### Oxidative stress

The Bos1-Sak1 signal transduction pathway was shown to be involved in oxidative stress response (Liu *et al.*, 2008). We based our analysis of proteins linked to oxidative stress (Table 3) on a recent review of Siegmund and Viefhues (Siegmund & Viefhues, 2016). Superoxide dismutase is involved in the detoxification process of superoxide ions. The deletion of *sak1* causes a decrease in the abundance of BcSod1 protein compared to the wild-type. Hydrogen peroxide is detoxified by the action of catalases. Among the eight catalases of the *B. cinerea* genome, three were found differentially produced in *Δsak1* and/or *Δbos1*. The Cat7 protein showed reduced abundance in the *Δsak1* mutant, while CatA was more abundant in the *Δbos1* mutant, compared to the wild-type. Catalase Cat5 is less present at both in *Δsak1* and *Δbos1.* Detoxification of ROS also requires the action of thioredoxins and disulfide-isomerases or peroxiredoxins (PRX). We observed a decrease in the biosynthesis of thioredoxin BcTRX1 and three disulfide isomerases in the *Δsak1* mutant. The abundance of only one disulfide isomerase was impacted in both *Δsak1* and *Δbos1* mutants. Out of nine PRX of the *B. cinerea* genome, three were identified as differentially abundant in *Δbos1* and *Δsak1* compared to the wild strain. PRX1 and PRX9 are less abundant in both *Δbos1* and *Δsak1.* In contrast, PRX8 is more abundant in both mutants.

Non-enzymatic mechanisms like the glutathione system are also involved in ROS detoxification. We found that four glutathione S-transferases (GST) were differentially abundant in *Δsak1* compared to the wild-type. GST 2 and 9 were less abundant while GST 5 and 14 were more abundant than the wild-type. Glutathione reductase BcGlr1 is important for the reduction of glutathione disulfide (GSSG) and the regeneration of glutathione (GSH). The abundance of this protein was decreased in the *Δbos1* mutant as well as in *Δsak1.* Glutoredoxin (Grx) is an oxidoreductase that is reduced via glutathione. Glutathione is then oxidized to GSSG. In the *Δsak1* mutant, the amount of glutoredoxin is lower compared to the wild-type.

Altogether, these data reveal important modifications in the content of proteins that play essential roles in ROS detoxification function in both ST mutants.

#### Cell wall stress

Fungal cell walls are mainly composed of chitin and β-1,3-glucans (Gow *et al.*, 2017). *Δbos1* and *Δsak1* mutants were shown to be affected in cell wall integrity (Liu *et al.*, 2011). Our proteomic study showed that in the *Δsak1* mutant, the abundance of both chitin synthases BcCHSIIIa and BcCHSV was increased compared to the wild type, while in the mutant *Δbos1* a decrease in the abundance of β-1,3-glucan synthase was observed.

### Crosslink between Sak1 and other signalling pathway proteins

The proteome of the *Δsak1* mutant was enriched in signalling proteins (Table 4). In particular, an accumulation of proteins involved in the G protein and cAMP pathways was observed. One of the three Gα subunits (BCG1) of the heterotrimeric complex Gαβγ was found overproduced in the *Δsak1* mutant. To a lesser extent, the CAP1 protein associated with adenylate cyclase and the regulatory subunit of protein kinase A (PKAR) were found more abundant. Thus, Sak1 negatively regulates the abundance of these proteins. In the *Δbos1* mutant, no difference in abundance these proteins compared to the wild-type was detected.

**Table 4:**
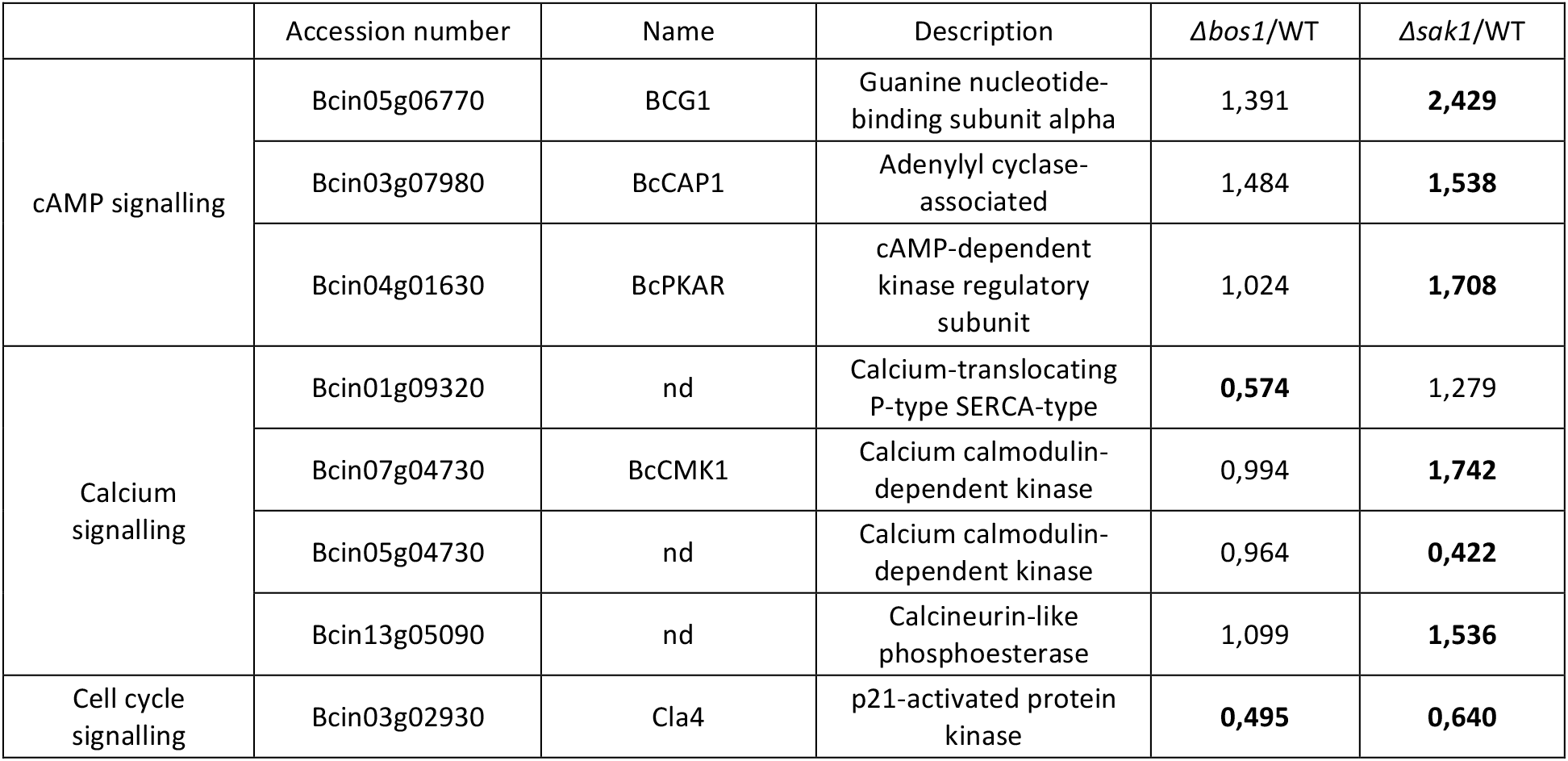
Signalling proteins of the AMPc, calcium and cell cycle signalling pathways whose abundance differs significantly in *Δbos1* and/or *Δsak1* mutants compared to wild type. Bold numbers indicate over threshold values.

| | Accession number | Name | Description | $\Delta bos1$/WT | $\Delta sak1$/WT |
| --- | --- | --- | --- | --- | --- |
| cAMP signalling | Bcin05g06770 | BCG1 | Guanine nucleotide-binding subunit alpha | 1,391 | **2,429** |
|  | Bcin03g07980 | BcCAP1 | Adenylyl cyclase-associated | 1,484 | **1,538** |
|  | Bcin04g01630 | BcPKAR | cAMP-dependent kinase regulatory subunit | 1,024 | **1,708** |
| Calcium signalling | Bcin01g09320 | nd | Calcium-translocating P-type SERCA-type | **0,574** | 1,279 |
|  | Bcin07g04730 | BcCMK1 | Calcium calmodulin-dependent kinase | 0,994 | **1,742** |
|  | Bcin05g04730 | nd | Calcium calmodulin-dependent kinase | 0,964 | **0,422** |
|  | Bcin13g05090 | nd | Calcineurin-like phosphoesterase | 1,099 | **1,536** |
| Cell cycle signalling | Bcin03g02930 | Cla4 | p21-activated protein kinase | **0,495** | **0,640** |

In order to confirm the detected link of Sak1 in cAMP pathway, we measured the cAMP concentration in both osmosensing mutants, *Δbos1* and *Δsak1.* In the *Δsak1* mutant, cAMP concentration was two to three-times higher compared to the wild-type strain (Figure 2). In contrast, the cAMP concentration in the *Δbos1* mutant was comparable to the wild-type confirming the negative regulation of the cAMP pathway by Sak1 independently of Bos1.

**Figure 2:**
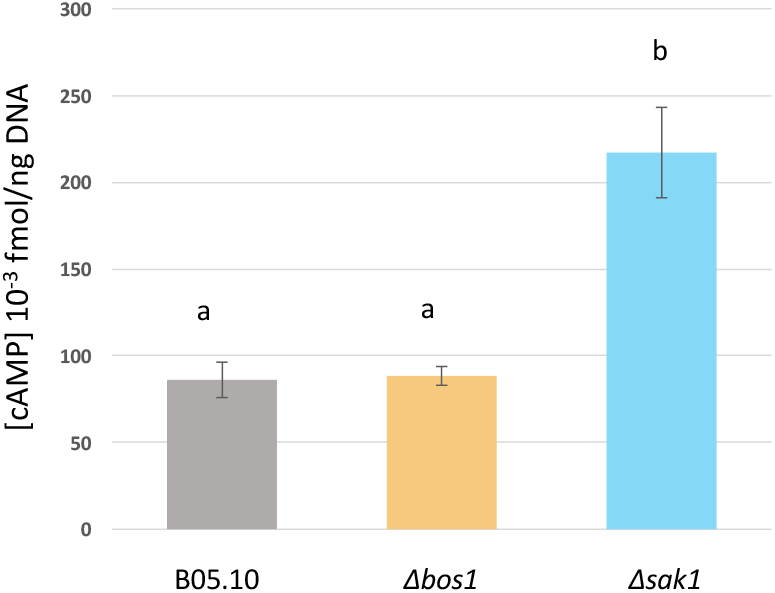
Intracellular concentration of cAMP in signal transduction mutants and parental strain B05.10. Values and standard deviations of three independent assays are indicated. Different letters indicate significantly different intracellular cAMP accumulation (P < 0,05).

In the *Δsak1* mutant, several proteins involved in the calcium signalling pathway such as two calcium calmodulin-dependent kinases and one calcineurin-like phosphoesterase also emerged from our analysis (Table 4). Both calcium calmodulin-dependent kinases were found regulated in opposite ways by Sak1: BcCMK1, was more abundant while the abundance of the second one was decreased. In the same strain, the abundance of a calcineurin-like phosphoesterase was slightly increased. Considering the *Δbos1* mutant, the above cited proteins did not vary, but a SERCA (Sarco/Endoplasmic Reticulum Ca^2+^ APTase) type calcium transporter was detected as less abundant than in the wild type.

In conclusion, our data clearly show an interconnection between the osmoregulating pathway and other signalling pathways, notably with G protein, cAMP and calcium signal transduction.

### Regulation of secreted proteases and cell wall degrading enzymes

The infection process of *B. cinerea,* especially early stages of infection, involves the secretion of plant cell wall degradating enzymes (CWDEs) (Table 5). The present proteomic analysis revealed 15 glycoside hydrolases, out of which 10 were less abundant in both *Δbos1* and *Δsak1* mutants. Moreover, two peptidases, one carboxypeptidase and two α/β-hydrolases were also less abundant in both mutants. Other enzymes, like one pectine lyase and one glycoside hydrolases were more abundant in *Δbos1* than in the wild-type while less abundant in *Δsak1.* Proteomic analysis also revealed an intracellular accumulation of CWDEs, annotated as secreted enzymes, especially in the *Δbos1* mutant. Among these proteins, glycoside hydrolases, pectin lyase A, pectate lyase A, endo-beta-1,4-glucanase Cel5A, and α/β- hydrolase were identified. We also noted the accumulation of endopolygalacturonase 1 BcPGA1 and a putative pectin methyltransferase. All these secreted proteins are involved in the modification and degradation of the plant wall and, consequently, in plant infection.

**Table 5:**
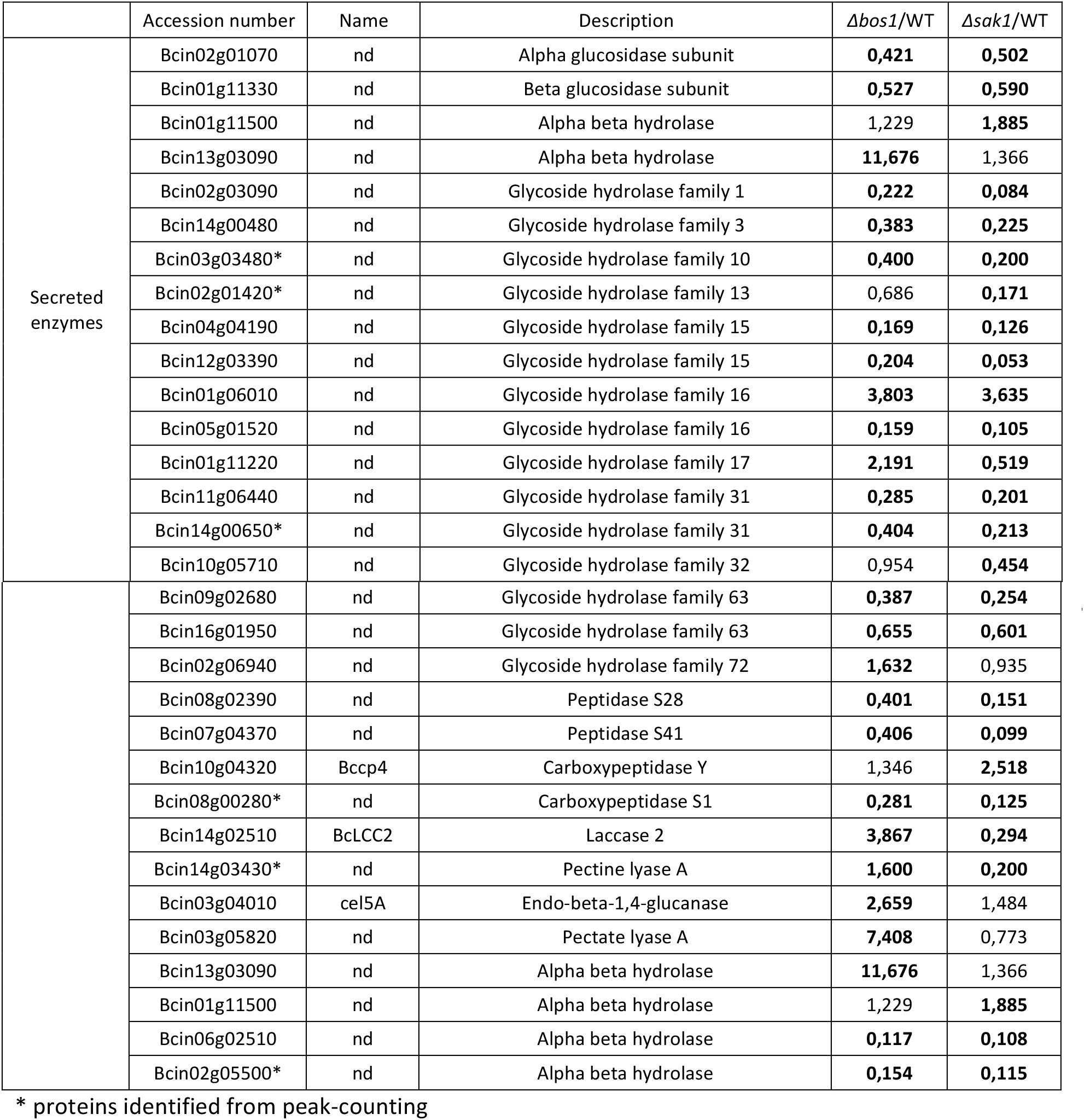
Putative secreted lytic enzymes, differentially produced in *Δbos1* and *Δsak1* mutants. Bold numbers indicate over threshold values.

| | Accession number | Name | Description | $\Delta bos1$/WT | $\Delta sak1$/WT |
| --- | --- | --- | --- | --- | --- |
| Secreted enzymes | Bcin02g01070 | nd | Alpha glucosidase subunit | **0,421** | **0,502** |
|  | Bcin01g11330 | nd | Beta glucosidase subunit | **0,527** | **0,590** |
|  | Bcin01g11500 | nd | Alpha beta hydrolase | 1,229 | **1,885** |
|  | Bcin13g03090 | nd | Alpha beta hydrolase | **11,676** | 1,366 |
|  | Bcin02g03090 | nd | Glycoside hydrolase family 1 | **0,222** | **0,084** |
|  | Bcin14g00480 | nd | Glycoside hydrolase family 3 | **0,383** | **0,225** |
|  | Bcin03g03480* | nd | Glycoside hydrolase family 10 | **0,400** | **0,200** |
|  | Bcin02g01420* | nd | Glycoside hydrolase family 13 | 0,686 | **0,171** |
|  | Bcin04g04190 | nd | Glycoside hydrolase family 15 | **0,169** | **0,126** |
|  | Bcin12g03390 | nd | Glycoside hydrolase family 15 | **0,204** | **0,053** |
|  | Bcin01g06010 | nd | Glycoside hydrolase family 16 | **3,803** | **3,635** |
|  | Bcin05g01520 | nd | Glycoside hydrolase family 16 | **0,159** | **0,105** |
|  | Bcin01g11220 | nd | Glycoside hydrolase family 17 | **2,191** | **0,519** |
|  | Bcin11g06440 | nd | Glycoside hydrolase family 31 | **0,285** | **0,201** |
|  | Bcin14g00650* | nd | Glycoside hydrolase family 31 | **0,404** | **0,213** |
|  | Bcin10g05710 | nd | Glycoside hydrolase family 32 | 0,954 | **0,454** |
|  | Bcin09g02680 | nd | Glycoside hydrolase family 63 | **0,387** | **0,254** |
|  | Bcin16g01950 | nd | Glycoside hydrolase family 63 | **0,655** | **0,601** |
|  | Bcin02g06940 | nd | Glycoside hydrolase family 72 | **1,632** | 0,935 |
|  | Bcin08g02390 | nd | Peptidase S28 | **0,401** | **0,151** |
|  | Bcin07g04370 | nd | Peptidase S41 | **0,406** | **0,099** |
|  | Bcin10g04320 | Bccp4 | Carboxypeptidase Y | 1,346 | **2,518** |
|  | Bcin08g00280* | nd | Carboxypeptidase S1 | **0,281** | **0,125** |
|  | Bcin14g02510 | BcLCC2 | Laccase 2 | **3,867** | **0,294** |
|  | Bcin14g03430* | nd | Pectine lyase A | **1,600** | **0,200** |
|  | Bcin03g04010 | cel5A | Endo-beta-1,4-glucanase | **2,659** | 1,484 |
|  | Bcin03g05820 | nd | Pectate lyase A | **7,408** | 0,773 |
|  | Bcin13g03090 | nd | Alpha beta hydrolase | **11,676** | 1,366 |
|  | Bcin01g11500 | nd | Alpha beta hydrolase | 1,229 | **1,885** |
|  | Bcin06g02510 | nd | Alpha beta hydrolase | **0,117** | **0,108** |
|  | Bcin02g05500* | nd | Alpha beta hydrolase | **0,154** | **0,115** |

\* proteins identified from peak-counting

## DISCUSSION

### Comparative proteomics of signal transduction mutants in Botrytis cinerea

In this work, we compared the steady-state protein levels of protein kinase mutants involved in osmosensing in the plant pathogenic fungus *Bortytis cinerea.* Analysing three different strains grown under axenic conditions led to the identification of roughly 1/5 of the predicted *B. cinerea* proteome. The abundance of 628 proteins (26% of the detected proteins) differed significantly in either or both mutants (>400 proteins per genotype). These results highlight the strong impact of protein kinase inactivation on protein production and turnover even in the absence of external stress. This is in agreement with other proteomic studies of fungal ST mutants (Zhang, H *et al.*, 2014; Isasa *et al.*, 2015; Liu *et al.*, 2018; Müller *et al.*, 2018) and, in the case of *Δsak1,* with the transcriptional results obtained by Heller and colleagues (Heller *et al.*, 2012).

### Regulation of protein abundance by Bos1 and/or Sak1

The protein kinases of this study are part of the same ST pathway. The histidine kinase Bos1 is a sensor-kinase of hyper-osmotic conditions (Viaud *et al.*, 2006) and negatively regulates the basal phosphorylation status of the Hog1-like MAPK Sak1 (Liu *et al.*, 2008). Comparing the proteomes of both mutants, we found that i/ the major part of the differential proteins (>60%) corresponds to proteins regulated independently by either Bos1 or Sak1, ii/ 43% of the differentially accumulating proteins are regulated by both protein kinases, but that iii/ only a small fraction (27 proteins) is controlled in opposing fashion by Bos1 and Sak1 which seems to be in contradiction to the negative regulation. Considering protein abundance – not the phosphorylation status – our results show that Bos1 and Sak1 act in a common direction under non-stressing conditions. The open question is: Might their opposing activities as suggested by (Liu *et al.*, 2008) have an impact only on the phosphorylation status of target proteins or could we have expected more contrasting proteomic data under stress conditions? One may expect contrasting impacts of *bos1* and *sak1* deletions on both, the phosphorylation status and protein abundance under stress conditions. In addition, the results obtained in this study also highlight a considerable part of independent functions for both kinases, which is in agreement with previous phenotypic studies (Segmuller *et al.*, 2007; Liu *et al.*, 2008; Liu *et al.*, 2011).

### Differentially abundant proteins are enriched in functional categories linked to infection

Functional enrichment studies of regulatory categories (proteins co-regulated or independently regulated by Bos1 and Sak1) revealed functions participating to the infection process and signalling proteins. Enzymes or proteins involved in secondary metabolism were enriched in most regulatory categories, revealing differential implications of the Bos1 and Sak1 kinases in their biosynthesis. Among the SM pathways detected, one can mention those of the toxins botrydial and botcinic acid, both involved in *B. cinerea* virulence (Dalmais *et al.*, 2011; Rossi *et al.*, 2011), but also oxalic acid. Although not essential to *B. cinerea* virulence, this organic acid contributes to lesion expansion through acidification essential for lytic enzymes (Kunz *et al.*, 2006; Müller *et al.*, 2018; Yin *et al.*, 2018). The strong reduction (100x) of oxaloacetate hydrolase (OAH) in the *Δsak1* mutant may certainly contribute to its reduced aggressiveness.

Among the proteins involved in stress resistance, our proteomic data revealed important changes for proteins involved in ROS production or detoxification. Especially, the *Δsak1* mutant had strongly reduced quantities of several ROS detoxifying enzymes (SOD, disulfide-isomerase, catalases). These are rather intriguing results. Even though both mutants are hypersensitive to hydrogen-peroxide and paraquat, sensitivity is even stronger in the *Δbos1* mutant. But this latter mutant also shows increased tolerance to menadione compared to the wild-type and the *Δsak1* mutant (Liu *et al.*, 2008). These phenotypes are not reflected by the differential steady-state levels of ROS detoxifying enzymes. We hypothesize that the constitutive production of the corresponding enzymes is not sufficient to prime the fungus against oxidative stress, but requires induced production under oxidative stress conditions controlled not only by the MAPK Sak1, but also by the MAPK Bmp3 and other yet unknown kinases, as highlighted by transcription analyses (Heller *et al.*, 2012).

A previous study showed the implication of Bos1 and Sak1 in cell wall integrity in *B. cinerea* without precising the respective cell wall modifications. Both *Δbos1* and *Δsak1* mutant are more sensitive than wild-type in regard to the two dyes, Congo red and Calcofluor White, but display differential sensitivity to other cell wall inhibitors (Liu *et al.*, 2011). The observed increase of chitine-synthase proteins (CHS3, CHS5) in the *Δsak1* mutant correlates nicely with its increased tolerance to nikkomycin Z, a specific inhibitor of *Saccharomyces cerevisiae* Chs3 (Gaughran *et al.*, 1994), and the related compound polyoxin B. The *Δbos1* mutant displays increased sensitivity to congo-red (Liu *et al.*, 2011), a component interacting with the glucan network of the fungal cell wall (Kopecká & Gabriel, 1992). This phenotype can be correlated to the decreased amount of beta-glucan-synthase observed for this mutant. Still, precise determination of cell wall composition is required to understand the mutants’ cell wall defects.

Among the enriched category of secreted lytic enzymes, we observed reductions in the large family of glycosyl-hydrolases in both mutants. In contrast, other lytic enzymes, especially plant cell wall degrading enzymes (CWDE) were found accumulating intracellularly in the *Δbos1* mutant. The underlying question is: Are these enzymes produced in higher amounts in this mutant or are they accumulating in the cytoplasm due to an eventual secretion defect? Part of the answer can be found considering proteins involved in secretion and vesicle transport. Some of them are less abundant in the *Δbos1* mutant (Table S1), *i.e.,* the Rab-GTPase Sas1, myosin, Sec18, Sey1, Yop1, Surf4, dynamin, all proteins involved in vesicle transport (Wang *et al.*, 1997; Steel *et al.*, 1999; Belden & Barlowe, 2001; Brands & Ho, 2002; Zhang, Z *et al.*, 2014), suggesting a potential secretion defect in the *Δbos1* mutant that may ultimately lead to the intracellular accumulation of plant CWDEs.

### Link to other signaling pathways

The here presented comparative proteomic data indicate a negative regulation of the cAMP pathway by the MAPK Sak1, reflected by the overproduction of one Ga sub-unit, Cap and PkaR proteins as well as of higher intracellular cAMP levels measured in the *Δsak1* mutant. An additional effect of the sak1-deletion was observed on proteins linked to Ca^2+^ signalling. This fits to the observation made by Schumacher and collaborators that the Ga sub-unit BCG1 is linked to the Ca^2^+ signalling pathway *via* PLC1 (Schumacher *et al.*, 2008), corroborating the links between the ST pathways identified by our proteomic data. These results also highlight that signalling proteins are not only regulated through phosphorylation and de-phosphorylation events, but also at the level of protein abundance. The relationship between the MAPK Sak1 the cAMP or Ca^2^+ signalling pathways may act at different levels: *via* phosphorylations and *via* the control of protein production and turnover.

Signalling pathways interact in all living organisms. In particular the hyperosmolarity and cell wall integrity pathways are known to interact in fungi (Fuchs & Mylonakis, 2009; Hamel *et al.*, 2012), as this was shown also for *B. cinerea* (Liu *et al.*, 2011; Heller *et al.*, 2012). Here we provide additional evidence for the involvement of the hyperosmolarity pathway in cell wall integrity, oxidative stress and its interaction with other signaling pathways. In addition, the observed changes in secondary metabolism proteins, notably those involved in botrytdial and botcinic acid biosynthesis corroborate interaction of Sak1 with the G-protein, cAMP and Ca^2+^ signaling. Indeed, all these pathways were shown to be involved in the regulation of transcription of BOT and BOA gene clusters and production of these SM (reviewed in (Viaud *et al.*, 2016)). The finding that OAH production is under Sak1 control is interesting as well. A recent publication revealed that OAH expression and the production of oxalate is controlled by the MAPKK of the CWI pathway *via* the protein kinase Sch9 without involvement of the MAPK Bmp3 (Yin *et al.*, 2018). It would now be interesting to analyse the respective roles of Sch9 and Sak1 in addition to their connection with respect to the regulation of OAH production.

In conclusion, proteomic analyses of ST mutants are complementary to phenotypic and transcriptomic analyses to characterize the biological functions of ST pathways, e.g., (Müller *et al.*, 2018). As the proteome reflects gene expression (i.e. transcription and RNA turn-over), protein production and turn-over, when combined with transcriptomics it may unravel additional regulations (e.g. protein degradation or secretion problems) of important function. Only few comparative proteomic studies of intracellular proteins of fungal ST mutants have been published so far (Zhang, H *et al.*, 2014; Isasa *et al.*, 2015; Liu *et al.*, 2018), although their potential to reveal regulations of unsuspected biological functions cannot be denied; e.g., the regulators of G-protein signalling proteins in *Magnaporthe oryzae* were shown to collectively regulate amino-acid metabolism (Zhang, H *et al.*, 2014), or, in the yeast *S. cerevisiae,* multiplex comparative proteomics highlighted implication of deubiquitylating proteins in mitochondrial regulation and phosphate metabolism (Isasa *et al.*, 2015). Ultimately, to dress the complete regulatory scheme of ST pathways and their downstream biological effects, time-lapse experiments including dynamic transcriptomic, proteomic and phosphoproteomic studies are required.

## ACKNOWLEDGEMENTS

The authors are grateful to region “Ile-de-France” and LABEX Saclay Plant Science (SPS) for financial support of proteomics equipment. They thank Adeline Simon for helping with enrichment studies.

## AUTHOR CONTRIBUTION

JK designed and performed all experiments, assisted by MD for the proteomics workflow. MZ performed proteomic data analysis and corresponding statistics. SF and MZ conceived the initial project. All authors participated to manuscript writing under the coordination of SF.

**Supporting Information Table S1:** List of differentially produced proteins in *Δbos1* and *Δsak1* mutants and fold-changes relative to wild-type. An asterisk follows the protein accession numbers identified by peak counting.

