## Supplemental Table S1 for "Comparative proteomics of osmotic signal transduction mutants in *Botrytis cinerea* explain loss of pathogenicity phenotypes and highlight interaction with cAMP and Ca^2+^ signalling pathways"

| **Bos1-** | **Accession number** | **Name** | **Description** | **Pfam** | ***Δbos1* / WT** | ***Δsak1* / WT** |
| --- | --- | --- | --- | --- | --- | --- |
|  | Bcin11g00330 | nd | hydroxymethylglutaryl- synthase | PF01154.12(HMG_CoA_synt_N) + PF08540.5(HMG_CoA_synt_C) | 1,513 | 0,697 |
|  | Bcin01g06440 | nd | translation initiation factor eIF3 subunit | PF08597.5(eIF3_subunit) | 1,532 | 1,119 |
|  | Bcin13g02240 | nd | importin beta-4 subunit | PF03810.14(IBN_N) + PF13513.1(HEAT_EZ) | 1,555 | 1,166 |
|  | Bcin04g05500 | nd | dihydroorotase | PF04909.9(Amidohydro_2) | 1,558 | 0,951 |
|  | Bcin05g08280 | nd | inosine-5 -monophosphate dehydrogenase imd2 | PF00478.20(IMPDH) + PF00571.23(CBS) + PF00571.23(CBS) | 1,561 | 1,017 |
|  | Bcin07g06150 | nd | BRO1-like domain-containing | PF03097.13(BRO1) + PF13949.1(ALIX_LYPXL_bnd) | 1,588 | 1,499 |
|  | Bcin12g04560 | BcGst24 | multisynthetase complex auxiliary component p43 | PF14497.1(GST_C_3) + PF01588.15(tRNA_bind) | 1,593 | 1,083 |
|  | Bcin07g01690 | nd | phosphoribosylaminoimidazole carboxylase | PF02222.17(ATP-grasp) + PF00731.15(AIRC) | 1,593 | 0,963 |
|  | Bcin07g05890 | nd | CTP synthase | PF06418.9(CTP_synth_N) + PF00117.23(GATase) | 1,595 | 1,314 |
|  | Bcin01g09170 | nd | branched-chain amino acid aminotransferase | PF01063.14(Aminotran_4) | 1,611 | 1,180 |
|  | Bcin13g04800 | nd | C2H2 and C2HC zinc finger | PF12756.2(zf-C2H2_2) + PF12756.2(zf-C2H2_2) | 1,613 | 1,229 |
|  | Bcin13g01760 | nd | nd | nd | 1,617 | 1,337 |
|  | Bcin14g04340 | nd | transcriptional co-repressor | PF00515.23(TPR_1) + PF13414.1(TPR_11) + PF00515.23(TPR_1) + PF00515.23(TPR_1) + PF00515.23(TPR_1) + PF13414.1(TPR_11) + PF07719.12(TPR_2) | 1,621 | 0,661 |
|  | Bcin03g01700 | nd | mitochondrial processing peptidase alpha subunit | PF00675.15(Peptidase_M16) + PF05193.16(Peptidase_M16_C) | 1,624 | 0,762 |
|  | Bcin02g02700 | nd | nd | nd | 1,628 | 1,490 |
|  | Bcin02g06940 | nd | glycoside hydrolase family 72 | PF03198.9(Glyco_hydro_72) + PF07983.8(X8) | 1,632 | 0,935 |
|  | Bcin15g04630 | nd | nd | PF00076.17(RRM_1) + PF00076.17(RRM_1) | 1,643 | 1,206 |
|  | Bcin10g03950 | nd | U3 small nucleolar RNA-associated 10 | PF12397.3(U3snoRNP10) + PF08146.7(BP28CT) | 1,648 | 0,874 |
|  | Bcin12g03930 | nd | 10 kDa heat shock mitochondrial | PF00166.16(Cpn10) | 1,656 | 1,300 |
|  | Bcin08g05590 | nd | WD40 repeat | PF00400.27(WD40) + PF00400.27(WD40) + PF00400.27(WD40) | 1,668 | 0,670 |
|  | Bcin16g04340 | nd | mitochondrial import receptor subunit TOM40 | PF01459.17(Porin_3) | 1,670 | 1,323 |
|  | Bcin02g06580 | nd | pyruvate decarboxylase | PF02776.13(TPP_enzyme_N) + PF00205.17(TPP_enzyme_M) + PF02775.16(TPP_enzyme_C) | 1,672 | 1,389 |
|  | Bcin15g01620 | nd | methylenetetrahydrofolate dehydrogenase | PF00763.18(THF_DHG_CYH) + PF02882.14(THF_DHG_CYH_C) | 1,683 | 1,245 |
|  | Bcin02g08410 | nd | mitochondrial glyco | PF02330.11(MAM33) | 1,691 | 0,867 |
|  | Bcin01g02090 | nd | phosphoribosylaminoimidazole-succinocarboxamide synthase | PF01259.13(SAICAR_synt) | 1,706 | 1,136 |
|  | Bcin07g04500 | nd | FK506-binding 4 | PF00254.23(FKBP_C) | 1,715 | 1,139 |
|  | Bcin09g02630 | nd | kh domain-containing | PF00013.24(KH_1) + PF00013.24(KH_1) + PF00013.24(KH_1) | 1,719 | 1,361 |
|  | Bcin05g02490 | nd | peptide methionine sulfoxide reductase | PF01625.16(PMSR) | 1,732 | 1,193 |
|  | Bcin07g01580 | nd | nucleolar GTP-binding | PF06858.9(NOG1) + PF08155.6(NOGCT) | 1,753 | 0,751 |
|  | Bcin02g05590 | nd | Ribosomal RNA small subunit methyltransferase NEP1 | PF03587.9(EMG1) | 1,756 | 0,844 |
|  | Bcin08g00910 | nd | glycosyl hydrolase family 17 | PF00332.13(Glyco_hydro_17) | 1,774 | 0,913 |
|  | Bcin07g00340 | nd | arginase | PF00491.16(Arginase) | 1,780 | 1,371 |
|  | Bcin01g01820 | nd | aspartate-semialdehyde dehydrogenase | PF01118.19(Semialdhyde_dh) + PF02774.13(Semialdhyde_dhC) | 1,787 | 1,388 |
|  | Bcin05g05580 | nd | ribose-phosphate pyrophosphokinase | PF13793.1(Pribosyltran_N) + PF14572.1(Pribosyl_synth) | 1,789 | 0,827 |
|  | Bcin06g02560 | nd | elongation factor mitochondrial | PF00009.22(GTP_EFTU) + PF03144.20(GTP_EFTU_D2) + PF14492.1(EFG_II) + PF03764.13(EFG_IV) + PF00679.19(EFG_C) | 1,801 | 1,408 |
|  | Bcin16g00830 | nd | Hsp90 co-chaperone Cdc37 | PF03234.9(CDC37_N) + PF08565.6(CDC37_M) + PF08564.5(CDC37_C) | 1,823 | 1,491 |
|  | Bcin01g10330 | nd | phospho-2-dehydro-3-deoxyheptonate aldolase | PF00793.15(DAHP_synth_1) | 1,825 | 1,462 |
|  | Bcin02g01760 | nd | phosphoribosylformylglycinamidine synthase | PF00586.19(AIRS) + PF02769.17(AIRS_C) + PF02769.17(AIRS_C) + PF13507.1(GATase_5) | 1,842 | 0,940 |
|  | Bcin03g05340 | nd | phosphoribosylamine-glycine ligase phosphoribosylformylglycinamidine cyclo-ligase | PF02844.10(GARS_N) + PF01071.14(GARS_A) + PF02843.11(GARS_C) + PF00586.19(AIRS) + PF02769.17(AIRS_C) | 1,859 | 1,213 |
|  | Bcin08g00630 | nd | glutamate-rich WD repeat containing 1 | PF12265.3(CAF1C_H4-bd) + PF00400.27(WD40) + PF00400.27(WD40) + PF00400.27(WD40) + PF00400.27(WD40) | 1,887 | 0,841 |
|  | Bcin06g03290 | nd | WD40 repeat | PF00400.27(WD40) + PF00400.27(WD40) + PF00400.27(WD40) | 1,897 | 0,928 |
|  | Bcin04g02970 | nd | chorismate synthase | PF01264.16(Chorismate_synt) | 1,910 | 1,176 |
|  | Bcin15g05360 | nd | galactose-1-phosphate uridylyltransferase | PF01087.17(GalP_UDP_transf) + PF02744.12(GalP_UDP_tr_C) | 1,934 | 0,935 |
|  | Bcin13g03880 | nd | adenylosuccinate lyase | PF00206.15(Lyase_1) + PF10397.4(ADSL_C) | 1,969 | 0,995 |
|  | Bcin11g00750 | nd | la domain family | PF05383.12(La) | 1,991 | 1,213 |
|  | Bcin09g02000 | nd | imidazole glycerol phosphate synthase hisHF | PF00117.23(GATase) + PF00977.16(His_biosynth) | 2,053 | 1,318 |
|  | Bcin13g04000 | nd | short-chain dehydrogenase | PF00106.20(adh_short) | 2,131 | 0,934 |
|  | Bcin10g04050 | nd | atp-dependent rna helicase drs1 | PF00270.24(DEAD) + PF00271.26(Helicase_C) | 2,224 | 1,074 |
|  | Bcin04g01520 | nd | Branched-chain-amino-acid aminotransferase | PF01063.14(Aminotran_4) | 2,477 | 1,377 |
|  | Bcin02g00280 | nd | fad binding domain | PF01565.18(FAD_binding_4) + PF08031.7(BBE) | 2,517 | 1,311 |
|  | Bcin03g04010 | cel5A | endo-beta-1,4-glucanase precursor | PF00150.13(Cellulase) | 2,659 | 1,484 |
|  | Bcin01g07240 | nd | DNA-binding HGH1 | PF04063.9(DUF383) + PF04064.8(DUF384) | 2,712 | 0,680 |
|  | Bcin12g01930 | nd | DNA-directed RNA polymerase I subunit | PF02150.11(RNA_POL_M_15KD) + PF01096.13(TFIIS_C) | 2,934 | 1,179 |
|  | Bcin07g00400 | nd | aldolase citrate lyase family | PF03328.9(HpcH_HpaI) | 3,011 | 0,998 |
|  | Bcin03g05820 | nd | pectate lyase a | PF00544.14(Pec_lyase_C) | 7,408 | 0,773 |
|  | Bcin13g04360 | nd | Aldo keto reductase | PF00248.16(Aldo_ket_red) | 9,213 | 0,689 |
|  | Bcin13g03090 | nd | alpha beta hydrolase fold | PF12697.2(Abhydrolase_6) | 11,676 | 1,366 |
|  | Bcin06g01180* | BcCatA | catalase | PF00199.14(Catalase) + PF06628.7(Catalase-rel) | 3,750 | 1,125 |

| **Bos1+** | **Accession number** | **Name** | **Description** | **Pfam** | ***Δbos1* / WT** | ***Δsak1* / WT** |
| --- | --- | --- | --- | --- | --- | --- |
|  | Bcin09g05010 | nd | fasciclin domain family | PF02469.17(Fasciclin) + PF02469.17(Fasciclin) | 0,102 | 0,957 |
|  | Bcin12g06380 | BcBot1 | cytochrome P450 | PF00067.17(p450) | 0,152 | 0,918 |
|  | Bcin08g07040 | nd | fatty acid hydroxylase | PF00067.17(p450) + PF00258.20(Flavodoxin_1) + PF00667.15(FAD_binding_1) + PF00175.16(NAD_binding_1) | 0,258 | 0,933 |
|  | Bcin04g05010 | nd | 3-oxo-5-alpha-steroid 4-dehydrogenase | PF02544.11(Steroid_dh) | 0,264 | 0,866 |
|  | Bcin09g06110 | nd | malate synthase | PF01274.17(Malate_synthase) | 0,273 | 0,678 |
|  | Bcin02g04510 | nd | nitroreductase | PF00881.19(Nitroreductase) | 0,368 | 1,024 |
|  | Bcin02g02320 | nd | oxidoreductase | PF01408.17(GFO_IDH_MocA) | 0,384 | 0,926 |
|  | Bcin02g02240 | nd | dolichyl-diphosphooligosaccharide- glycosyltransferase | PF02516.9(STT3) | 0,404 | 0,780 |
|  | Bcin12g00750 | nd | cation-transporting atpase 4 | PF00122.15(E1-E2_ATPase) + PF00702.21(Hydrolase) | 0,404 | 1,223 |
|  | Bcin05g07500 | nd | Amidase signature (AS) enzyme | PF01425.16(Amidase) | 0,410 | 0,745 |
|  | Bcin05g07200 | nd | nd | nd | 0,410 | 0,827 |
|  | Bcin09g01320 | nd | isocitrate lyase | PF00463.16(ICL) | 0,420 | 1,242 |
|  | Bcin16g02170 | nd | cleft lip and palate associated transmembrane 1 | PF05602.7(CLPTM1) | 0,422 | 0,776 |
|  | Bcin08g04420 | nd | IMP-specific 5 -nucleotidase 1 | PF06437.6(ISN1) | 0,426 | 1,222 |
|  | Bcin01g02310 | BcPmt1 | dolichyl-phosphate-mannose- mannosyltransferase | PF02366.13(PMT) + PF02815.14(MIR) | 0,430 | 1,141 |
|  | Bcin04g01430 | nd | nd | nd | 0,434 | 0,897 |
|  | Bcin12g05330 | nd | glycolipid transfer | PF08718.6(GLTP) | 0,456 | 0,783 |
|  | Bcin04g05050 | nd | e3 ubiquitin- ligase huwe1 | PF06012.7(DUF908) + PF06025.7(DUF913) + PF00627.26(UBA) + PF14377.1(DUF4414) + PF00632.20(HECT) | 0,462 | 0,728 |
|  | Bcin01g00550 | BcSas1 | GTP-binding ypt2 | PF00071.17(Ras) | 0,466 | 0,835 |
|  | Bcin01g07510 | nd | Uncharacterized J domain-containing | PF00226.26(DnaJ) | 0,479 | 1,052 |
|  | Bcin05g02240 | nd | bap31 domain-containing | PF05529.7(Bap31) | 0,481 | 0,955 |
|  | Bcin11g04180 | nd | surf4 family | PF02077.10(SURF4) | 0,487 | 0,818 |
|  | Bcin01g03500 | nd | C2 (Calcium lipid- ) | PF00168.25(C2) + PF00168.25(C2) + PF00168.25(C2) + PF00168.25(C2) + PF00168.25(C2) | 0,488 | 1,055 |
|  | Bcin02g06930 | nd | 1,3-beta-glucan synthase | PF14288.1(FKS1_dom1) + PF02364.10(Glucan_synthase) | 0,491 | 1,309 |
|  | Bcin15g02970 | nd | 5 -3 exoribonuclease 1 | PF03159.13(XRN_N) | 0,495 | 0,885 |
|  | Bcin11g06340 | nd | ubiquitin-conjugating enzyme | PF00179.21(UQ_con) | 0,495 | 0,779 |
|  | Bcin15g03420 | BcPmt2 | dolichyl-phosphate-mannose- mannosyltransferase | PF02366.13(PMT) + PF02815.14(MIR) | 0,498 | 1,139 |
|  | Bcin12g05090 | nd | vacuolar ATP synthase subunit d | PF01813.12(ATP-synt_D) | 0,500 | 0,979 |
|  | Bcin01g08400 | nd | oligosaccharyltransferase subunit ribophorin II | PF05817.9(Ribophorin_II) | 0,500 | 0,792 |
|  | Bcin08g05920 | nd | dynamin family | PF00350.18(Dynamin_N) + PF01031.15(Dynamin_M) | 0,501 | 0,696 |
|  | Bcin03g00430 | nd | heat shock 70 | PF00012.15(HSP70) + PF00012.15(HSP70) | 0,502 | 0,757 |
|  | Bcin07g03080 | nd | Metalloproteases (zincins) catalytic | PF01433.15(Peptidase_M1) + PF11838.3(ERAP1_C) | 0,503 | 0,996 |
|  | Bcin05g00650 | nd | allantoicase | PF03561.10(Allantoicase) + PF03561.10(Allantoicase) | 0,504 | 1,245 |
|  | Bcin06g06260 | nd | 20S proteasome subunit beta 2 | PF00227.21(Proteasome) + PF12465.3(Pr_beta_C) | 0,509 | 0,932 |
|  | Bcin02g06610 | nd | Chaperone J-domain-containing | PF13414.1(TPR_11) + PF13371.1(TPR_9) + PF13371.1(TPR_9) + PF00226.26(DnaJ) | 0,511 | 0,784 |
|  | Bcin03g02370 | nd | yop1 | PF03134.14(TB2_DP1_HVA22) | 0,520 | 1,227 |
|  | Bcin06g02430 | nd | aminotransferase class i and ii | PF00155.16(Aminotran_1_2) | 0,523 | 0,764 |
|  | Bcin01g03020 | nd | methylmalonate-semialdehyde dehydrogenase | PF00171.17(Aldedh) | 0,532 | 0,948 |
|  | Bcin16g03180 | nd | acyl- dehydrogenase family member 11 | PF02771.11(Acyl-CoA_dh_N) + PF02770.14(Acyl-CoA_dh_M) + PF00441.19(Acyl-CoA_dh_1) | 0,534 | 0,692 |
|  | Bcin06g04360 | nd | cop9 signalosome complex subunit 2 | PF01399.22(PCI) | 0,538 | 0,879 |
|  | Bcin02g01980 | BcPmt4 | dolichyl-phosphate-mannose- mannosyltransferase | PF02366.13(PMT) + PF02815.14(MIR) | 0,540 | 0,946 |
|  | Bcin14g01750 | nd | carbon-nitrogen hydrolase | PF00795.17(CN_hydrolase) | 0,541 | 0,988 |
|  | Bcin05g00360 | nd | nd | PF00063.16(Myosin_head) + PF06017.8(Myosin_TH1) + PF00018.23(SH3_1) | 0,544 | 1,239 |
|  | Bcin02g03080 | nd | succinate dehydrogenase subunit C | PF01127.17(Sdh_cyt) | 0,546 | 1,159 |
|  | Bcin12g02810 | nd | mannan polymerase ii complex anp1 subunit | PF03452.9(Anp1) | 0,548 | 0,988 |
|  | Bcin01g03640 | nd | UTP-glucose-1-phosphate uridylyltransferase | PF01704.13(UDPGP) | 0,549 | 0,785 |
|  | Bcin06g02540 | nd | rpel repeat | PF02755.10(RPEL) + PF02755.10(RPEL) | 0,562 | 1,182 |
|  | Bcin04g04420 | nd | transcription factor C2H2 | PF00627.26(UBA) | 0,571 | 0,991 |
|  | Bcin01g09320 | nd | calcium-translocating P-type SERCA-type | PF00690.21(Cation_ATPase_N) + PF00122.15(E1-E2_ATPase) + PF00702.21(Hydrolase) + PF00689.16(Cation_ATPase_C) | 0,574 | 1,279 |
|  | Bcin01g03230 | nd | prolyl oligopeptidase | PF00326.16(Peptidase_S9) | 0,575 | 0,768 |
|  | Bcin12g00320 | nd | glycolipid 2-alpha-mannosyltransferase | PF01793.11(Glyco_transf_15) | 0,578 | 0,669 |
|  | Bcin08g02840 | nd | dolichyl-diphosphooligosaccharide-- glycosyltransferase 48 kDa subunit | PF03345.9(DDOST_48kD) | 0,583 | 0,779 |
|  | Bcin11g04630 | nd | glyoxylate reductase | PF00389.25(2-Hacid_dh) + PF02826.14(2-Hacid_dh_C) | 0,586 | 1,205 |
|  | Bcin03g03150 | nd | 3-hydroxybutyryl- dehydrogenase | PF02737.13(3HCDH_N) + PF00725.17(3HCDH) | 0,589 | 0,925 |
|  | Bcin02g04240 | nd | fatty acid elongase 2 | PF01151.13(ELO) | 0,590 | 1,480 |
|  | Bcin08g04730 | nd | fructose-1,6-bisphosphatase | PF00316.15(FBPase) | 0,590 | 0,815 |
|  | Bcin04g01150 | nd | OST3 OST6 family | PF04756.8(OST3_OST6) | 0,590 | 0,877 |
|  | Bcin05g00940 | nd | cyclopropane-fatty-acyl-phospholipid synthase | PF02353.15(CMAS) | 0,593 | 1,051 |
|  | Bcin03g05840 | nd | enoyl- hydratase | PF00378.15(ECH) | 0,596 | 0,718 |
|  | Bcin15g00500 | nd | nadh-ubiquinone oxidoreductase subunit grim-19 | PF06212.7(GRIM-19) | 0,597 | 1,000 |
|  | Bcin15g03930 | nd | endoplasmic reticulum | PF03661.8(UPF0121) | 0,600 | 1,006 |
|  | Bcin01g01700 | nd | phosphoinositide phosphatase | PF02383.13(Syja_N) | 0,601 | 1,204 |
|  | Bcin15g04980 | nd | AGC AKT kinase | PF00069.20(Pkinase) + PF00433.19(Pkinase_C) | 0,601 | 0,775 |
|  | Bcin11g06270 | nd | delta-1-pyrroline-5-carboxylate dehydrogenase | PF00171.17(Aldedh) | 0,615 | 0,669 |
|  | Bcin07g02610 | nd | 6-phosphogluconate dehydrogenase | PF03446.10(NAD_binding_2) + PF00393.14(6PGD) | 0,618 | 0,877 |
|  | Bcin02g00160 | nd | cipC-like antibiotic response | PF12585.3(DUF3759) | 0,618 | 0,916 |
|  | Bcin10g05230 | nd | 4-aminobutyrate aminotransferase | PF00202.16(Aminotran_3) | 0,618 | 1,202 |
|  | Bcin02g02740 | nd | mitochondrial DNA replication YHM2 | PF00153.22(Mito_carr) + PF00153.22(Mito_carr) + PF00153.22(Mito_carr) | 0,621 | 1,151 |
|  | Bcin15g02610 | nd | cell division control 12 | PF00735.13(Septin) | 0,621 | 0,981 |
|  | Bcin07g01610 | nd | aspartyl protease | PF00240.18(ubiquitin) + PF09668.5(Asp_protease) + PF00627.26(UBA) | 0,622 | 1,160 |
|  | Bcin03g07910 | nd | carnitine O-acetyltransferase | PF00755.15(Carn_acyltransf) | 0,622 | 1,184 |
|  | Bcin14g00830 | nd | PLP-dependent transferase | PF00266.14(Aminotran_5) | 0,629 | 1,496 |
|  | Bcin11g03990 | nd | nd | PF11489.3(DUF3210) | 0,632 | 0,690 |
|  | Bcin07g02010 | nd | nd | PF12701.2(LSM14) + PF09532.5(FDF) | 0,633 | 0,757 |
|  | Bcin16g03680 | nd | P-loop containing nucleoside triphosphate hydrolase | PF09439.5(SRPRB) | 0,635 | 1,011 |
|  | Bcin07g02150 | nd | histone h2a | PF00125.19(Histone) | 0,636 | 0,907 |
|  | Bcin07g00840 | nd | peptidyl-prolyl cis-trans isomerase B | PF00160.16(Pro_isomerase) | 0,638 | 0,761 |
|  | Bcin12g03270 | nd | Zn-dependent exopeptidase | PF04389.12(Peptidase_M28) | 0,643 | 0,696 |
|  | Bcin03g06080 | nd | class V myosin (Myo4) | PF00063.16(Myosin_head) + PF00612.22(IQ) + PF00612.22(IQ) + PF00612.22(IQ) + PF01843.14(DIL) | 0,646 | 0,965 |
|  | Bcin05g05700 | nd | vesicular-fusion sec18 | PF02359.13(CDC48_N) + PF02933.12(CDC48_2) + PF00004.24(AAA) + PF00004.24(AAA) | 0,647 | 0,706 |
|  | Bcin15g01350 | nd | root hair defective 3 GTP-binding | PF05879.7(RHD3) | 0,647 | 1,274 |
|  | Bcin11g05210 | nd | nmda receptor-regulated 1 | PF00515.23(TPR_1) + PF12569.3(NARP1) | 0,648 | 0,935 |
|  | Bcin11g01880 | nd | membrane-associated progesterone receptor component 1 | PF00173.23(Cyt-b5) | 0,648 | 1,292 |
|  | Bcin01g09510 | nd | proteasome component c5 | PF00227.21(Proteasome) | 0,654 | 0,931 |
|  | Bcin03g02440 | nd | nd | PF13233.1(Complex1_LYR_2) | 0,654 | 0,838 |
|  | Bcin02g08340 | BcTps1 | glycosyltransferase family 20 | PF00982.16(Glyco_transf_20) | 0,657 | 0,722 |
|  | Bcin11g02490 | nd | phosphoglucomutase | PF02878.11(PGM_PMM_I) + PF02879.11(PGM_PMM_II) + PF02880.11(PGM_PMM_III) + PF00408.15(PGM_PMM_IV) | 0,657 | 0,761 |
|  | Bcin11g01660 | nd | inosine-uridine preferring nucleoside hydrolase | PF01156.14(IU_nuc_hydro) | 0,579 | 0,664 |
|  | Bcin15g00860 | nd | prenylcysteine oxidase | PF13450.1(NAD_binding_8) + PF07156.9(Prenylcys_lyase) | 0,529 | 0,662 |

| **Sak1-** | **Accession number** | **Name** | **Description** | **Pfam** | ***Δbos1* / WT** | ***Δsak1* / WT** |
| --- | --- | --- | --- | --- | --- | --- |
|  | Bcin08g03470 | nd | DUF1620 domain-containing | PF13360.1(PQQ_2) + PF07774.8(DUF1620) | 1,017 | 1,506 |
|  | Bcin09g06820 | nd | ubx domain | PF14555.1(UBA_4) + PF00789.15(UBX) | 1,146 | 1,507 |
|  | Bcin03g06600 | nd | hsp70 | PF00012.15(HSP70) | 1,324 | 1,507 |
|  | Bcin09g03820 | nd | sh3 domain-containing | PF00018.23(SH3_1) | 1,397 | 1,507 |
|  | Bcin01g06220 | nd | gtp-binding nuclear gsp1 ran | PF00071.17(Ras) | 1,455 | 1,507 |
|  | Bcin10g02440 | nd | Cytosolic J-domain-containing | PF13414.1(TPR_11) + PF13174.1(TPR_6) + PF07719.12(TPR_2) + PF13414.1(TPR_11) + PF00226.26(DnaJ) | 1,252 | 1,521 |
|  | Bcin02g02540 | nd | mitochondrial carrier | PF00153.22(Mito_carr) + PF00153.22(Mito_carr) | 1,190 | 1,521 |
|  | Bcin13g05020 | nd | threonine synthase | PF14821.1(Thr_synth_N) + PF00291.20(PALP) | 1,285 | 1,522 |
|  | Bcin08g04540 | nd | pyruvate dehydrogenase E1 component beta subunit | PF02779.19(Transket_pyr) + PF02780.15(Transketolase_C) | 1,349 | 1,528 |
|  | Bcin09g00600 | nd | short-chain dehydrogenase | PF00106.20(adh_short) | 1,297 | 1,528 |
|  | Bcin04g02200 | nd | acetolactate synthase small subunit | PF01842.20(ACT) + PF10369.4(ALS_ss_C) | 1,262 | 1,536 |
|  | Bcin13g05090 | nd | calcineurin-like phosphoesterase | PF00149.23(Metallophos) | 1,099 | 1,536 |
|  | Bcin03g07980 | nd | adenylyl cyclase-associated | PF01213.14(CAP_N) + PF08603.6(CAP_C) | 1,484 | 1,538 |
|  | Bcin04g04490 | nd | asparagine synthetase | PF13537.1(GATase_7) + PF00733.16(Asn_synthase) | 1,421 | 1,541 |
|  | Bcin03g06180 | nd | D-3-phosphoglycerate dehydrogenase | PF00389.25(2-Hacid_dh) + PF02826.14(2-Hacid_dh_C) | 1,285 | 1,542 |
|  | Bcin05g05970 | nd | vacuolar atp synthase 98 kda subunit | PF01496.14(V_ATPase_I) | 0,859 | 1,543 |
|  | Bcin09g04160 | nd | hsp70 | PF00012.15(HSP70) | 0,912 | 1,552 |
|  | Bcin04g03430 | nd | CK1 CK1 CK1-G kinase | PF00069.20(Pkinase) | 0,896 | 1,554 |
|  | Bcin14g04870 | nd | phb1 prohibitin 1 | PF01145.20(Band_7) | 1,245 | 1,557 |
|  | Bcin03g01890 | nd | S-phase kinase-associated 1 | PF03931.10(Skp1_POZ) + PF01466.14(Skp1) | 1,065 | 1,572 |
|  | Bcin02g00470 | nd | L-xylulose reductase | PF13561.1(adh_short_C2) | 0,742 | 1,576 |
|  | Bcin11g03430 | nd | ubiquitin SMT3 | PF11976.3(Rad60-SLD) | 1,499 | 1,577 |
|  | Bcin08g03520 | nd | alkaline phosphatase | PF00245.15(Alk_phosphatase) | 1,128 | 1,585 |
|  | Bcin01g00220 | nd | Altered inheritance of mitochondria mitochondrial | PF04588.8(HIG_1_N) | 1,417 | 1,586 |
|  | Bcin04g03120 | BcChsIIIa | chitin synthase g | PF08407.6(Chitin_synth_1N) + PF01644.12(Chitin_synth_1) | 0,697 | 1,591 |
|  | Bcin15g00320 | nd | hypothetical protein BC1G_12400 | PF11563.3(Protoglobin) | 1,283 | 1,595 |
|  | Bcin05g04270 | nd | saccharopine dehydrogenase | PF05222.10(AlaDh_PNT_N) + PF01262.16(AlaDh_PNT_C) | 1,330 | 1,615 |
|  | Bcin05g00010 | nd | Aldo keto reductase | PF00248.16(Aldo_ket_red) | 0,927 | 1,616 |
|  | Bcin12g00930 | nd | carbohydrate-binding module family 50 | PF05433.10(Rick_17kDa_Anti) + PF08881.5(CVNH) + PF01476.15(LysM) | 0,924 | 1,617 |
|  | Bcin05g02400 | nd | mitochondrial hypoxia responsive domain-containing | PF04588.8(HIG_1_N) | 1,439 | 1,623 |
|  | Bcin02g02890 | nd | nucleoporin nic96 | PF13634.1(Nucleoporin_FG) + PF13634.1(Nucleoporin_FG) + PF13634.1(Nucleoporin_FG) + PF04097.9(Nic96) | 0,826 | 1,627 |
|  | Bcin01g06330 | nd | pyridoxal kinase | PF08543.7(Phos_pyr_kin) | 1,487 | 1,628 |
|  | Bcin12g05370 | BcChsV | chitin synthase | PF00063.16(Myosin_head) + PF00173.23(Cyt-b5) + PF00173.23(Cyt-b5) + PF03142.10(Chitin_synth_2) + PF08766.6(DEK_C) | 0,736 | 1,638 |
|  | Bcin13g02990 | nd | bsd domain-containing | PF03909.12(BSD) | 1,278 | 1,646 |
|  | Bcin09g06240 | nd | WH1 domain-containing | PF00568.18(WH1) | 1,167 | 1,650 |
|  | Bcin02g01850 | nd | PX domain-containing | PF12828.2(PXB) + PF00787.19(PX) + PF12825.2(DUF3818) | 0,676 | 1,658 |
|  | Bcin11g04890 | nd | glycine cleavage system t | PF01571.16(GCV_T) + PF08669.6(GCV_T_C) | 1,055 | 1,668 |
|  | Bcin14g04420 | nd | nd | PF14388.1(DUF4419) | 0,858 | 1,673 |
|  | Bcin13g04010 | nd | sh3 domain-containing | PF04366.7(DUF500) + PF07653.12(SH3_2) | 1,139 | 1,678 |
|  | Bcin02g07550 | nd | cbs pb1 domain-containing | PF00571.23(CBS) + PF00571.23(CBS) + PF00571.23(CBS) + PF00571.23(CBS) + PF00564.19(PB1) | 1,436 | 1,704 |
|  | Bcin14g02370 | BcCyc1 | cytochrome c | PF00034.16(Cytochrom_C) | 1,407 | 1,707 |
|  | Bcin09g00980 | nd | isocitrate dehydrogenase | PF00180.15(Iso_dh) | 0,910 | 1,708 |
|  | Bcin04g01630 | BcPkar | camp-dependent kinase regulatory subunit | PF02197.12(RIIa) + PF00027.24(cNMP_binding) + PF00027.24(cNMP_binding) | 1,024 | 1,708 |
|  | Bcin04g01050 | nd | Calponin- CH-domain-containing | PF13499.1(EF-hand_7) + PF00307.26(CH) + PF00307.26(CH) + PF00307.26(CH) + PF00307.26(CH) | 1,151 | 1,720 |
|  | Bcin05g03270 | nd | mrs7 family | PF07766.8(LETM1) | 1,262 | 1,730 |
|  | Bcin06g07240 | nd | MC family mitochondrial carrier | PF00153.22(Mito_carr) + PF00153.22(Mito_carr) + PF00153.22(Mito_carr) | 0,847 | 1,732 |
|  | Bcin09g07030 | nd | glucosamine-phosphate N-acetyltransferase | PF13508.1(Acetyltransf_7) | 1,187 | 1,732 |
|  | Bcin15g02860 | nd | mitochondrial ribosomal | PF01245.15(Ribosomal_L19) | 0,901 | 1,740 |
|  | Bcin07g04730 | BcCmk1 | calcium calmodulin-dependent kinase | PF00069.20(Pkinase) | 0,994 | 1,742 |
|  | Bcin16g02680 | nd | peptidase m16 inactive domain-containing | PF00675.15(Peptidase_M16) + PF05193.16(Peptidase_M16_C) + PF05193.16(Peptidase_M16_C) | 1,333 | 1,742 |
|  | Bcin02g00910 | nd | uba ts-n domain containing | PF12763.2(EF-hand_4) + PF12763.2(EF-hand_4) + PF00627.26(UBA) | 1,266 | 1,745 |
|  | Bcin14g04940 | nd | aminotransferase class i and ii | PF00155.16(Aminotran_1_2) | 1,119 | 1,747 |
|  | Bcin01g06930 | BcCarA | carbamoyl-phosphate synthase subunit arginine-specific small | PF08252.6(Leader_CPA1) | 1,320 | 1,750 |
|  | Bcin10g03910 | nd | hsp98 | PF02861.15(Clp_N) + PF02861.15(Clp_N) + PF00004.24(AAA) + PF07724.9(AAA_2) + PF10431.4(ClpB_D2-small) | 1,073 | 1,752 |
|  | Bcin02g00170 | nd | Vacuolar aminopeptidase 1 | PF02127.10(Peptidase_M18) | 1,284 | 1,758 |
|  | Bcin11g01920 | nd | vacuolar sorting-associated 35 | PF03635.12(Vps35) | 0,761 | 1,774 |
|  | Bcin16g02820 | nd | dihydroxy-acid dehydratase | PF00920.16(ILVD_EDD) | 1,211 | 1,781 |
|  | Bcin04g02090 | nd | ubiquitin elongating factor core | PF10408.4(Ufd2P_core) + PF04564.10(U-box) | 0,972 | 1,794 |
|  | Bcin14g02120 | nd | CUE domain-containing | PF02845.11(CUE) | 1,113 | 1,800 |
|  | Bcin13g01610 | nd | alanine transaminase | PF00155.16(Aminotran_1_2) | 1,349 | 1,826 |
|  | Bcin10g02660 | nd | hypothetical protein BC1G_04690 | PF04828.9(GFA) | 1,337 | 1,837 |
|  | Bcin01g03110 | nd | pre-mRNA-processing factor 39 | nd | 0,671 | 1,841 |
|  | Bcin11g00970 | nd | branched-chain-amino-acid aminotransferase | PF01063.14(Aminotran_4) | 1,246 | 1,844 |
|  | Bcin01g02160 | nd | cytoskeleton assembly control Sla2 | PF07651.11(ANTH) + PF01608.12(I_LWEQ) | 1,265 | 1,859 |
|  | Bcin02g01720 | nd | phospholipase D nuclease | PF00614.17(PLDc) + PF00614.17(PLDc) | 0,764 | 1,859 |
|  | Bcin04g03100 | nd | Ketol-acid mitochondrial | PF07991.7(IlvN) + PF01450.14(IlvC) | 1,407 | 1,866 |
|  | Bcin14g02720 | nd | cysteine desulfurase | PF00266.14(Aminotran_5) | 1,108 | 1,869 |
|  | Bcin07g04330 | nd | exportin 1 | PF03810.14(IBN_N) + PF08389.7(Xpo1) | 0,834 | 1,870 |
|  | Bcin15g05110 | nd | 2-nitropropane dioxygenase | PF03060.10(NMO) | 1,373 | 1,872 |
|  | Bcin01g11500 | nd | alpha beta hydrolase | PF12697.2(Abhydrolase_6) | 1,229 | 1,885 |
|  | Bcin07g06160 | nd | atp-dependent protease la | PF02190.11(LON) + PF00004.24(AAA) + PF05362.8(Lon_C) | 1,429 | 1,901 |
|  | Bcin04g01680 | nd | iron sulfur cluster assembly 1 | PF01592.11(NifU_N) | 1,296 | 1,910 |
|  | Bcin12g05290 | nd | glutaminyl-tRNA synthetase | PF00749.16(tRNA-synt_1c) + PF03950.13(tRNA-synt_1c_C) | 0,856 | 1,956 |
|  | Bcin01g07090 | nd | ferric-chelate reductase | PF01794.14(Ferric_reduct) + PF08022.7(FAD_binding_8) + PF08030.7(NAD_binding_6) | 1,124 | 1,992 |
|  | Bcin09g04730 | nd | E1 -activating enzyme G | PF00899.16(ThiF) | 1,456 | 1,995 |
|  | Bcin04g06230 | BcFhg1 | flavohemo | PF00042.17(Globin) + PF00970.19(FAD_binding_6) + PF00175.16(NAD_binding_1) | 1,441 | 2,001 |
|  | Bcin08g00720 | BcAp14 | pepsinogen c | PF00026.18(Asp) | 1,304 | 2,010 |
|  | Bcin11g04370 | nd | ethanolamine kinase | PF01633.15(Choline_kinase) | 1,443 | 2,023 |
|  | Bcin11g03040 | nd | acetolactate synthase | PF02776.13(TPP_enzyme_N) + PF00205.17(TPP_enzyme_M) + PF02775.16(TPP_enzyme_C) | 1,302 | 2,063 |
|  | Bcin08g00090 | nd | sulfide quinone reductase | PF07992.9(Pyr_redox_2) | 1,437 | 2,083 |
|  | Bcin16g01000 | CND8 | p-loop containing nucleoside triphosphate hydrolase | PF00004.24(AAA) | 1,166 | 2,083 |
|  | Bcin07g06310 | nd | acyl- desaturase | PF00487.19(FA_desaturase) + PF00173.23(Cyt-b5) | 1,130 | 2,090 |
|  | Bcin06g00800 | nd | inositol monophosphatase | PF00459.20(Inositol_P) | 1,484 | 2,150 |
|  | Bcin12g01280 | nd | sh3 domain-containing | PF07653.12(SH3_2) | 1,049 | 2,221 |
|  | Bcin08g01460 | nd | glycine dehydrogenase | PF02347.11(GDC-P) + PF02347.11(GDC-P) | 1,186 | 2,225 |
|  | Bcin06g05570 | nd | duf1772 domain containing | PF08592.6(DUF1772) | 0,781 | 2,242 |
|  | Bcin16g00630 | nd | phosphoenolpyruvate carboxykinase | PF01293.15(PEPCK_ATP) | 0,896 | 2,245 |
|  | Bcin07g04130 | nd | short chain dehydrogenase | PF00106.20(adh_short) + PF00106.20(adh_short) + PF13452.1(MaoC_dehydrat_N) + PF01575.14(MaoC_dehydratas) | 0,799 | 2,307 |
|  | Bcin12g03740 | nd | sulfite reductase hemo beta-component | PF01855.14(POR_N) + PF00258.20(Flavodoxin_1) + PF03460.12(NIR_SIR_ferr) + PF01077.17(NIR_SIR) + PF03460.12(NIR_SIR_ferr) | 1,218 | 2,350 |
|  | Bcin03g05100 | nd | glycerol-3-phosphate o-acyltransferase | PF01553.16(Acyltransferase) | 0,917 | 2,392 |
|  | Bcin12g04570 | nd | phosphomevalonate kinase | PF00288.21(GHMP_kinases_N) + PF08544.8(GHMP_kinases_C) | 1,176 | 2,429 |
|  | Bcin05g06770 | BcBcg1 | Guanine nucleotide-binding subunit alpha | PF00503.15(G-alpha) | 1,391 | 2,429 |
|  | Bcin10g04320 | BcCp4 | carboxypeptidase y | PF05388.6(Carbpep_Y_N) + PF00450.17(Peptidase_S10) | 1,346 | 2,518 |
|  | Bcin14g02730 | nd | reduced viability upon starvation | PF03114.13(BAR) + PF00018.23(SH3_1) | 1,141 | 2,524 |
|  | Bcin05g06820 | nd | 3-hydroxyacyl- dehydrogenase | PF00106.20(adh_short) | 1,250 | 2,633 |
|  | Bcin06g05690 | nd | proline iminopeptidase | PF00561.15(Abhydrolase_1) | 0,940 | 2,728 |
|  | Bcin10g02060 | nd | Uncharacterized oxidoreductase yvaA | PF01408.17(GFO_IDH_MocA) + PF02894.12(GFO_IDH_MocA_C) | 0,777 | 2,737 |
|  | Bcin15g00990 | nd | spindle poison sensitivity Scp3 | PF00642.19(zf-CCCH) | 0,864 | 2,742 |
|  | Bcin08g04440 | nd | ser Thr phosphatase family | PF00149.23(Metallophos) | 0,888 | 2,816 |
|  | Bcin03g04100 | nd | nd | PF07217.6(Het-C) | 1,460 | 2,830 |
|  | Bcin08g03270 | nd | vacuolar 8 | PF00514.18(Arm) + PF00514.18(Arm) + PF00514.18(Arm) + PF00514.18(Arm) + PF00514.18(Arm) + PF00514.18(Arm) + PF00514.18(Arm) | 1,471 | 2,877 |
|  | Bcin14g03790 | nd | sphingosine-1-phosphate lyase | PF00282.14(Pyridoxal_deC) | 0,951 | 2,919 |
|  | Bcin01g05510 | nd | CTR2 long splice variant | PF04145.10(Ctr) | 1,243 | 3,042 |
|  | Bcin01g00160 | BcBoA17 | 3-oxoacyl-[acyl-carrier- ] reductase | PF00106.20(adh_short) | 0,965 | 3,202 |
|  | Bcin13g05210 | nd | high affinity copper transporter | PF04145.10(Ctr) | 1,110 | 3,222 |
|  | Bcin15g04430 | nd | methyltransferase domain-containing | PF08241.7(Methyltransf_11) | 1,357 | 3,327 |
|  | Bcin07g06550 | nd | 3-isopropylmalate dehydratase | PF00330.15(Aconitase) + PF00694.14(Aconitase_C) | 1,140 | 3,447 |
|  | Bcin13g01710 | nd | FAD-binding monooxygenase | PF01494.14(FAD_binding_3) | 0,990 | 3,514 |
|  | Bcin10g05310 | nd | 3-isopropylmalate dehydrogenase | PF00180.15(Iso_dh) | 1,428 | 3,554 |
|  | Bcin01g08020 | nd | D-isomer-specific 2-hydroxyacid dehydrogenase | PF02826.14(2-Hacid_dh_C) + PF02826.14(2-Hacid_dh_C) | 1,389 | 3,672 |
|  | Bcin03g06830 | nd | uricase | PF01014.13(Uricase) + PF01014.13(Uricase) | 1,054 | 3,679 |
|  | Bcin10g00060 | nd | oxidoreductase NAD-binding domain-containing | PF00970.19(FAD_binding_6) + PF00175.16(NAD_binding_1) | 1,162 | 3,784 |
|  | Bcin07g03430 | BcGst14 | glutathione S-transferase domain-containing | PF13417.1(GST_N_3) | 0,762 | 3,980 |
|  | Bcin03g03940 | nd | 3-ketoacyl-coa thiolase peroxisomal a precursor | PF00108.18(Thiolase_N) + PF02803.13(Thiolase_C) | 1,065 | 4,187 |
|  | Bcin01g11550 | BcPks5 | polyketide synthase | PF00109.21(ketoacyl-synt) + PF02801.17(Ketoacyl-synt_C) + PF00698.16(Acyl_transf_1) + PF14765.1(PS-DH) + PF08242.7(Methyltransf_12) + PF08659.5(KR) + PF00668.15(Condensation) + PF13745.1(HxxPF_rpt) + PF00501.23(AMP-binding) + PF00550.20(PP-binding) + PF07993.7(NAD_binding_4) | 0,843 | 4,753 |
|  | Bcin01g11520 | nd | non-ribosomal peptide synthetase | nd | 1,401 | 9,058 |
|  | Bcin01g11470 | nd | nd | nd | 1,130 | 12,699 |
|  | Bcin01g11490 | nd | O- family 2 | PF00891.13(Methyltransf_2) | 1,286 | 15,438 |
|  | Bcin08g01120* | nd | bleomycin hydrolase | PF03051.10(Peptidase_C1_2) | 1,107 | 1,714 |
|  | Bcin16g00570* | nd | DHS-like NAD FAD-binding domain-containing | PF02146.12(SIR2) | 1,320 | 2,320 |
|  | Bcin16g03120* | nd | nadp-dependent malic enzyme | PF00390.14(malic) + PF03949.10(Malic_M) | 1,375 | 2,458 |
|  | Bcin02g07090* | nd | NADH:flavin oxidoreductase NADH oxidase | PF00724.15(Oxidored_FMN) | 1,143 | 2,714 |
|  | Bcin01g11480* | nd | bifunctional p-450:nadph-p450 reductase | PF00067.17(p450) + PF00258.20(Flavodoxin_1) + PF00667.15(FAD_binding_1) + PF00175.16(NAD_binding_1) | 1,306 | 4,000 |

| **Sak1+** | **Accession number** | **Name** | **Description** | **Pfam** | ***Δbos1* / WT** | ***Δsak1* / WT** |
| --- | --- | --- | --- | --- | --- | --- |
|  | Bcin01g05720 | nd | extracellular serine-rich | nd | 1,220 | 0,055 |
|  | Bcin04g04800 | BcBrn1 | tetrahydroxynaphthalene reductase | PF00106.20(adh_short) | 1,434 | 0,088 |
|  | Bcin02g00230 | nd | alcohol dehydrogenase | PF00248.16(Aldo_ket_red) | 1,099 | 0,097 |
|  | Bcin02g00270 | nd | nd | PF07110.6(EthD) | 0,921 | 0,168 |
|  | Bcin10g01780 | nd | nd | PF14021.1(DUF4237) | 0,927 | 0,195 |
|  | Bcin12g02050 | nd | alcohol dehydrogenase | PF08240.7(ADH_N) + PF00107.21(ADH_zinc_N) | 0,867 | 0,204 |
|  | Bcin07g06060 | nd | short-chain dehydrogenase reductase SDR | PF00106.20(adh_short) | 1,420 | 0,255 |
|  | Bcin14g03970 | nd | 1,3-beta-glucanosyltransferase gel1 | PF03198.9(Glyco_hydro_72) | 1,484 | 0,266 |
|  | Bcin03g03390 | BcSod1 | superoxide dismutase | PF00080.15(Sod_Cu) | 0,727 | 0,283 |
|  | Bcin05g07670 | nd | alcohol dehydrogenase | PF08240.7(ADH_N) + PF00107.21(ADH_zinc_N) | 0,723 | 0,303 |
|  | Bcin11g04060 | nd | nd | nd | 0,899 | 0,327 |
|  | Bcin01g06800 | nd | phospho-2-dehydro-3-deoxyheptonate aldolase | PF01474.11(DAHP_synth_2) | 1,000 | 0,330 |
|  | Bcin12g01010 | nd | nd | nd | 0,703 | 0,342 |
|  | Bcin12g02450 | nd | biotin synthase | PF04055.16(Radical_SAM) + PF06968.8(BATS) | 0,675 | 0,350 |
|  | Bcin05g00670 | nd | peptidase family M13 | PF05649.8(Peptidase_M13_N) + PF01431.16(Peptidase_M13) | 0,735 | 0,353 |
|  | Bcin03g04790 | nd | aldo keto reductase | PF00248.16(Aldo_ket_red) | 1,119 | 0,374 |
|  | Bcin10g02280 | nd | glycosyl transferase family 8 | PF01501.15(Glyco_transf_8) | 0,829 | 0,381 |
|  | Bcin15g02270 | nd | glycogen synthase | PF05693.8(Glycogen_syn) | 0,695 | 0,385 |
|  | Bcin08g05050 | nd | isovaleryl- dehydrogenase | PF02771.11(Acyl-CoA_dh_N) + PF02770.14(Acyl-CoA_dh_M) + PF00441.19(Acyl-CoA_dh_1) | 0,752 | 0,401 |
|  | Bcin05g04730 | nd | calcium calmodulin-dependent kinase | PF00069.20(Pkinase) | 0,964 | 0,422 |
|  | Bcin16g04350 | nd | elongator complex 2 | PF01591.13(6PF2K) + PF00300.17(His_Phos_1) | 0,697 | 0,447 |
|  | Bcin06g03360 | nd | nd | nd | 1,122 | 0,452 |
|  | Bcin10g05710 | nd | glycoside hydrolase family 32 | PF00251.15(Glyco_hydro_32N) | 0,954 | 0,454 |
|  | Bcin09g00280 | nd | zinc knuckle domain | PF00098.18(zf-CCHC) + PF00098.18(zf-CCHC) + PF00098.18(zf-CCHC) + PF00098.18(zf-CCHC) + PF00098.18(zf-CCHC) + PF00098.18(zf-CCHC) + PF14392.1(zf-CCHC_4) | 1,008 | 0,467 |
|  | Bcin02g02110 | nd | ribulose-phosphate 3-epimerase | PF00834.14(Ribul_P_3_epim) | 0,912 | 0,478 |
|  | Bcin03g09270 | nd | phosphatase 5 | PF13414.1(TPR_11) + PF00515.23(TPR_1) + PF08321.7(PPP5) + PF00149.23(Metallophos) | 0,671 | 0,480 |
|  | Bcin11g04640 | nd | anthranilate phosphoribosyltransferase | PF02885.12(Glycos_trans_3N) + PF00591.16(Glycos_transf_3) | 0,994 | 0,495 |
|  | Bcin12g04280 | BcTrx1 | thioredoxin | PF00085.15(Thioredoxin) | 0,814 | 0,496 |
|  | Bcin01g10500 | nd | nd | PF08881.5(CVNH) | 0,761 | 0,498 |
|  | Bcin02g01020 | nd | conserved fungal | PF07958.6(DUF1688) | 0,828 | 0,499 |
|  | Bcin16g03380 | nd | mannitol dehydrogenase | PF00106.20(adh_short) | 0,837 | 0,501 |
|  | Bcin03g05330 | nd | pyridoxine biosynthesis | PF01680.12(SOR_SNZ) | 0,726 | 0,509 |
|  | Bcin03g03060 | nd | dTDP-glucose 4,6-dehydratase | PF01370.16(Epimerase) | 1,077 | 0,509 |
|  | Bcin10g03330 | nd | mrna processing | PF13824.1(zf-Mss51) | 1,106 | 0,514 |
|  | Bcin01g09450 | BcLgd1 | l-galactonate dehydratase | PF01188.16(MR_MLE) + PF13378.1(MR_MLE_C) | 0,993 | 0,525 |
|  | Bcin05g02250 | nd | tRNA-dihydrouridine synthase 3 | PF01207.12(Dus) + PF01207.12(Dus) | 1,056 | 0,527 |
|  | Bcin09g03020 | nd | histone h1-binding | PF10516.4(SHNi-TPR) | 0,737 | 0,543 |
|  | Bcin04g06240 | nd | 2-nitropropane dioxygenase | PF03060.10(NMO) | 0,719 | 0,543 |
|  | Bcin07g03460 | nd | survival factor 1 | PF08622.5(Svf1) | 0,663 | 0,552 |
|  | Bcin01g06070 | nd | eukaryotic phosphomannomutase | PF03332.8(PMM) | 1,028 | 0,555 |
|  | Bcin03g07940 | nd | 60s ribosomal l3 | nd | 1,223 | 0,559 |
|  | Bcin06g05730 | BcPdi1 | disulfide isomerase | PF00085.15(Thioredoxin) + PF13848.1(Thioredoxin_6) + PF00085.15(Thioredoxin) | 0,753 | 0,573 |
|  | Bcin07g04070 | nd | nd | nd | 1,217 | 0,578 |
|  | Bcin03g04170 | nd | imidazoleglycerol-phosphate dehydratase | PF00475.13(IGPD) | 1,124 | 0,579 |
|  | Bcin12g00280 | nd | phosphoserine phosphatase | PF12710.2(HAD) | 1,074 | 0,583 |
|  | Bcin02g03060 | PpoA80 | fatty acid oxygenase | PF03098.10(An_peroxidase) + PF00067.17(p450) | 1,066 | 0,590 |
|  | Bcin16g03500 | nd | cystathionine beta-synthase | PF00291.20(PALP) + PF00571.23(CBS) | 1,242 | 0,591 |
|  | Bcin12g01360 | BcPIC5, BcFKBP12 | peptidylprolyl isomerase | PF00254.23(FKBP_C) | 0,828 | 0,595 |
|  | Bcin06g01640 | nd | succinate semialdehyde dehydrogenase | PF00171.17(Aldedh) | 0,933 | 0,599 |
|  | Bcin07g04620 | nd | f-actin capping beta subunit | PF01115.12(F_actin_cap_B) | 0,736 | 0,602 |
|  | Bcin09g02240 | nd | oxidoreductase family | PF01408.17(GFO_IDH_MocA) | 0,985 | 0,604 |
|  | Bcin05g00380 | nd | nd | nd | 0,721 | 0,606 |
|  | Bcin02g08570 | nd | sec14 cytosolic factor | PF03765.10(CRAL_TRIO_N) + PF00650.15(CRAL_TRIO) | 0,897 | 0,615 |
|  | Bcin11g05910 | nd | Dipeptidyl-peptidase III | PF03571.10(Peptidase_M49) | 0,671 | 0,627 |
|  | Bcin02g06270 | nd | 2,3-bisphosphoglycerate-independent phosphoglycerate mutase | PF01676.13(Metalloenzyme) + PF06415.8(iPGM_N) | 1,073 | 0,629 |
|  | Bcin06g05400 | nd | acetyl- acetyltransferase | PF00108.18(Thiolase_N) + PF02803.13(Thiolase_C) | 0,755 | 0,629 |
|  | Bcin04g00340 | BcPtc3 | phosphatase 2C | PF00481.16(PP2C) | 0,804 | 0,632 |
|  | Bcin01g08280 | nd | phosphoribosylglycinamide formyltransferase | PF00551.14(Formyl_trans_N) | 1,294 | 0,634 |
|  | Bcin10g01240 | BcGrx1 | glutaredoxin | PF00462.19(Glutaredoxin) | 0,750 | 0,635 |
|  | Bcin15g02120 | nd | glyceraldehyde-3-phosphate dehydrogenase | PF00044.19(Gp_dh_N) + PF02800.15(Gp_dh_C) | 0,939 | 0,635 |
|  | Bcin01g08850 | nd | S-methyl-5-thioribose-1-phosphate isomerase | PF01008.12(IF-2B) | 1,021 | 0,636 |
|  | Bcin06g01850 | nd | zinc finger gcs1 | PF01412.13(ArfGap) | 0,969 | 0,639 |
|  | Bcin05g00580 | nd | UDP-galactopyranose mutase | PF13450.1(NAD_binding_8) | 0,702 | 0,640 |
|  | Bcin09g05650 | nd | nd | nd | 0,914 | 0,644 |
|  | Bcin01g08260 | nd | 6,7-dimethyl-8-ribityllumazine synthase | PF00885.14(DMRL_synthase) + PF00885.14(DMRL_synthase) | 0,818 | 0,651 |
|  | Bcin01g10660 | nd | transporter sec-13 | PF00400.27(WD40) + PF00400.27(WD40) + PF00400.27(WD40) + PF00400.27(WD40) + PF00400.27(WD40) + PF00400.27(WD40) | 0,886 | 0,651 |
|  | Bcin03g07930 | nd | La domain-containing | PF05383.12(La) + PF14259.1(RRM_6) | 1,153 | 0,652 |
|  | Bcin05g08370 | nd | disulfide isomerase | PF00085.15(Thioredoxin) | 0,827 | 0,653 |
|  | Bcin09g01850 | nd | duf1687 domain containing | PF07955.6(DUF1687) | 0,768 | 0,657 |
|  | Bcin02g05070 | nd | Sec23 Sec24 family | PF04810.10(zf-Sec23_Sec24) + PF04811.10(Sec23_trunk) + PF08033.7(Sec23_BS) + PF04815.10(Sec23_helical) + PF00626.17(Gelsolin) | 1,196 | 0,659 |
|  | Bcin09g04400* | BcCat7 | catalase domain containing | PF00199.14(Catalase) | 1,040 | 0,040 |
|  | Bcin03g08110* | BcScd1 | conidial pigment biosynthesis scytalone dehydratase Arp1 | PF02982.9(Scytalone_dh) | 0,826 | 0,043 |
|  | Bcin01g04900* | BcGstII | glutathione s-transferase | PF02798.15(GST_N) + PF00043.20(GST_C) | 1,125 | 0,156 |
|  | Bcin05g07190* | nd | Class I glutamine amidotransferase | PF13587.1(DJ-1_PfpI_N) + PF01965.19(DJ-1_PfpI) | 1,320 | 0,160 |
|  | Bcin02g01420* | nd | glycoside hydrolase family 13 | PF00128.19(Alpha-amylase) + PF09260.6(DUF1966) | 0,686 | 0,171 |
|  | Bcin03g01570* | nd | NRPS-like enzyme | PF00501.23(AMP-binding) + PF00550.20(PP-binding) + PF07993.7(NAD_binding_4) | 1,077 | 0,192 |
|  | Bcin05g02050* | BcAlo1 | AMID-like mitochondrial oxidoreductase | PF07992.9(Pyr_redox_2) + PF00070.22(Pyr_redox) | 1,000 | 0,205 |
|  | Bcin08g05060* | nd | nad-dependent 15-hydroxyprostaglandin dehydrogenase | nd | 0,667 | 0,212 |
|  | Bcin06g01930* | BcGo1 | carbohydrate-Binding Module family 18 | PF07250.6(Glyoxal_oxid_N) + PF09118.6(DUF1929) | 1,067 | 0,244 |
|  | Bcin08g02950* | nd | D-arabinitol dehydrogenase | PF13561.1(adh_short_C2) | 0,794 | 0,265 |
|  | Bcin15g05600* | CND9 | mannitol-1-phosphate 5-dehydrogenase | PF01232.18(Mannitol_dh) + PF08125.8(Mannitol_dh_C) | 0,767 | 0,333 |
|  | Bcin07g02980* | nd | NADH:flavin oxidoreductase NADH oxidase family | PF00724.15(Oxidored_FMN) | 1,269 | 0,346 |
|  | Bcin16g04990* | nd | nd | nd | 1,174 | 0,348 |
|  | Bcin07g01720* | nd | oligopeptidase family | PF00326.16(Peptidase_S9) | 1,146 | 0,375 |
|  | Bcin12g02040* | BcAp8 | aspartic protease | PF00026.18(Asp) | 0,733 | 0,378 |
|  | Bcin07g06070* | nd | acyl- dehydrogenase domain-containing | PF02771.11(Acyl-CoA_dh_N) + PF02770.14(Acyl-CoA_dh_M) + PF00441.19(Acyl-CoA_dh_1) | 1,034 | 0,414 |
|  | Bcin10g00140* | nd | Short-chain dehydrogenase reductase SDR | PF00106.20(adh_short) | 1,185 | 0,444 |
|  | Bcin02g03910* | nd | aldehyde dehydrogenase | PF00171.17(Aldedh) | 0,853 | 0,500 |

| **Bos1- / Sak1-** | **Accession number** | **Name** | **Description** | **Pfam** | ***Δbos1* / WT** | ***Δsak1* / WT** |
| --- | --- | --- | --- | --- | --- | --- |
|  | Bcin16g00820 | nd | atp-dependent permease mdl2 | PF00664.18(ABC_membrane) + PF00005.22(ABC_tran) | 1,527 | 2,158 |
|  | Bcin09g02500 | nd | 60s ribosomal l44 | nd | 1,530 | 1,954 |
|  | Bcin15g03910 | nd | palmitoyl- thioesterase precursor | PF02089.10(Palm_thioest) | 1,533 | 3,417 |
|  | Bcin02g01780 | nd | trans-aconitate 2-methyltransferase | PF08241.7(Methyltransf_11) | 1,538 | 2,088 |
|  | Bcin08g04430 | nd | o-methyltransferase family | PF01596.12(Methyltransf_3) | 1,552 | 2,974 |
|  | Bcin01g01990 | nd | rna binding | PF00076.17(RRM_1) | 1,557 | 1,757 |
|  | Bcin01g08220 | nd | zinc-binding oxidoreductase | PF08240.7(ADH_N) + PF00107.21(ADH_zinc_N) | 1,561 | 2,764 |
|  | Bcin09g02210 | nd | adenylosuccinate synthetase | PF00709.16(Adenylsucc_synt) | 1,565 | 1,563 |
|  | Bcin09g05610 | nd | nd | nd | 1,575 | 1,938 |
|  | Bcin10g00300 | nd | heat shock 90 | PF13589.1(HATPase_c_3) + PF00183.13(HSP90) | 1,592 | 1,509 |
|  | Bcin14g00430 | nd | homoaconitase | PF00330.15(Aconitase) + PF00330.15(Aconitase) + PF00694.14(Aconitase_C) | 1,599 | 1,830 |
|  | Bcin16g02120 | nd | CCR4-Not complex subunit Caf16 | PF00005.22(ABC_tran) | 1,601 | 1,641 |
|  | Bcin03g02170 | nd | mitochondrial inner membrane AAA protease Yta12 | PF06480.10(FtsH_ext) + PF00004.24(AAA) + PF01434.13(Peptidase_M41) | 1,603 | 2,066 |
|  | Bcin12g00640 | nd | nd | nd | 1,627 | 4,022 |
|  | Bcin07g04780 | nd | PLP-dependent transferase | PF00155.16(Aminotran_1_2) | 1,629 | 3,911 |
|  | Bcin07g00730 | nd | ornithine aminotransferase | PF00202.16(Aminotran_3) | 1,635 | 1,678 |
|  | Bcin06g07090 | nd | ATP-dependent RNA helicase | PF00270.24(DEAD) + PF00271.26(Helicase_C) | 1,641 | 1,547 |
|  | Bcin01g09530 | nd | nd | PF00011.16(HSP20) | 1,658 | 5,466 |
|  | Bcin05g00690 | nd | nd | nd | 1,662 | 1,853 |
|  | Bcin02g00540 | nd | prolyl oligopeptidase | PF07859.8(Abhydrolase_3) | 1,687 | 2,129 |
|  | Bcin08g03710 | BcNde2 | NADH dehydrogenase | PF07992.9(Pyr_redox_2) + PF00070.22(Pyr_redox) | 1,688 | 2,004 |
|  | Bcin02g03340 | nd | ran gtpase activating 1 | PF13516.1(LRR_6) | 1,726 | 1,757 |
|  | Bcin07g06220 | nd | heat shock 70 kDa | PF00012.15(HSP70) | 1,729 | 1,550 |
|  | Bcin03g09280 | nd | saccharopine dehydrogenase | PF03435.13(Saccharop_dh) | 1,730 | 2,495 |
|  | Bcin08g03960 | nd | nd | nd | 1,737 | 2,097 |
|  | Bcin06g02300 | nd | homoisocitrate dehydrogenase | PF00180.15(Iso_dh) | 1,741 | 2,819 |
|  | Bcin11g02580 | nd | Mitochondrial import inner membrane translocase subunit tim44 | PF04280.10(Tim44) | 1,755 | 1,596 |
|  | Bcin06g06290 | nd | cytosine deaminase | PF00383.17(dCMP_cyt_deam_1) | 1,789 | 6,283 |
|  | Bcin13g03450 | nd | sulfite reductase flavo component | PF01558.13(POR) + PF00667.15(FAD_binding_1) + PF00175.16(NAD_binding_1) | 1,806 | 2,527 |
|  | Bcin13g01330 | nd | 5 -methylthioadenosine phosphorylase | PF01048.15(PNP_UDP_1) | 1,809 | 1,770 |
|  | Bcin13g04430 | nd | pentafunctional arom polypeptide | PF01761.15(DHQ_synthase) + PF00275.15(EPSP_synthase) + PF01202.17(SKI) + PF01487.10(DHquinase_I) + PF08501.6(Shikimate_dh_N) + PF01488.15(Shikimate_DH) | 1,854 | 1,899 |
|  | Bcin16g02010 | nd | hsp70 | PF08609.5(Fes1) + PF13513.1(HEAT_EZ) | 1,868 | 1,774 |
|  | Bcin12g05000 | nd | chorismate mutase | nd | 1,869 | 2,440 |
|  | Bcin04g00140 | nd | l-aminoadipate-semialdehyde dehydrogenase large subunit | PF00501.23(AMP-binding) + PF13193.1(AMP-binding_C) + PF00550.20(PP-binding) + PF07993.7(NAD_binding_4) | 1,875 | 2,602 |
|  | Bcin02g06470 | BcStr2 | cystathionine gamma-synthase | PF01053.15(Cys_Met_Meta_PP) | 1,919 | 1,933 |
|  | Bcin01g11530 | nd | zinc-binding dehydrogenase family | PF00107.21(ADH_zinc_N) | 1,929 | 19,547 |
|  | Bcin09g02600 | nd | like mitochondrial | PF01025.14(GrpE) | 1,929 | 1,698 |
|  | Bcin09g00730 | nd | siderophore iron transporter mirB | PF07690.11(MFS_1) | 1,942 | 3,732 |
|  | Bcin14g00540 | nd | myo-inositol-1-phosphate synthase | PF07994.7(NAD_binding_5) + PF01658.12(Inos-1-P_synth) | 1,961 | 1,527 |
|  | Bcin07g05720 | nd | ubiquitin carboxyl-terminal hydrolase | PF06337.7(DUSP) + PF00443.24(UCH) | 1,978 | 9,411 |
|  | Bcin01g01190 | BcAp3 | aspartic-type endopeptidase | PF00026.18(Asp) | 1,985 | 2,054 |
|  | Bcin01g04340 | nd | sulfate adenylyltransferase | PF14306.1(PUA_2) + PF01747.12(ATP-sulfurylase) + PF01583.15(APS_kinase) | 1,985 | 2,390 |
|  | Bcin09g03930 | BcPrx8 | peroxisomal matrix | PF08534.5(Redoxin) | 2,011 | 7,046 |
|  | Bcin14g04440 | nd | nd | nd | 2,025 | 7,328 |
|  | Bcin02g08210 | nd | ngg1 interacting factor | PF01784.13(NIF3) | 2,097 | 1,666 |
|  | Bcin04g06830 | nd | nd | nd | 2,147 | 1,519 |
|  | Bcin03g04900 | nd | hypothetical protein BC1G_00781 | PF05022.7(SRP40_C) | 2,194 | 1,734 |
|  | Bcin08g05470 | nd | choline-sulfatase | PF00884.18(Sulfatase) + PF12411.3(Choline_sulf_C) | 2,204 | 6,363 |
|  | Bcin14g01480 | nd | dsba oxidoreductase | PF01323.15(DSBA) | 2,216 | 1,935 |
|  | Bcin11g05230 | nd | mitochondrial import inner membrane translocase subunit tim54 | PF11711.3(Tim54) | 2,224 | 1,967 |
|  | Bcin02g02580 | CPD2 | coproporphyrinogen iii oxidase | PF01218.13(Coprogen_oxidas) | 2,231 | 3,906 |
|  | Bcin11g03020 | nd | galactokinase | PF10509.4(GalKase_gal_bdg) + PF00288.21(GHMP_kinases_N) + PF08544.8(GHMP_kinases_C) | 2,246 | 1,519 |
|  | Bcin04g06310 | nd | Metal homeostasis factor atx1 | PF00403.21(HMA) | 2,303 | 3,276 |
|  | Bcin14g05400 | nd | homocitrate synthase | PF00682.14(HMGL-like) | 2,325 | 2,550 |
|  | Bcin15g01020 | nd | chitin synthase activator | PF08238.7(Sel1) + PF08238.7(Sel1) + PF08238.7(Sel1) + PF08238.7(Sel1) + PF08238.7(Sel1) | 2,351 | 1,732 |
|  | Bcin09g06840 | nd | nd | PF14033.1(DUF4246) | 2,387 | 14,999 |
|  | Bcin01g00210 | nd | threonine ammonia- biosynthetic | PF00291.20(PALP) + PF00585.13(Thr_dehydrat_C) + PF00585.13(Thr_dehydrat_C) | 2,395 | 2,513 |
|  | Bcin09g03710 | nd | amidophosphoribosyltransferase | PF00310.16(GATase_2) + PF00156.22(Pribosyltran) | 2,415 | 1,504 |
|  | Bcin01g01040 | nd | WD40 repeat | PF00400.27(WD40) | 2,443 | 1,891 |
|  | Bcin15g02310 | nd | nd | PF05368.8(NmrA) | 2,493 | 2,568 |
|  | Bcin01g05550 | nd | carbamoyl-phosphate large subunit | PF00289.17(CPSase_L_chain) + PF02786.12(CPSase_L_D2) + PF02787.14(CPSase_L_D3) + PF00289.17(CPSase_L_chain) + PF02786.12(CPSase_L_D2) | 2,575 | 2,097 |
|  | Bcin03g05720 | nd | nd | nd | 2,582 | 7,796 |
|  | Bcin14g00860 | B431 | carbohydrate esterase family 8 | PF01095.14(Pectinesterase) | 2,618 | 2,727 |
|  | Bcin02g05740 | nd | allantoinase | PF01979.15(Amidohydro_1) | 2,656 | 5,639 |
|  | Bcin06g02010 | nd | Copper-binding of the mitochondrial inner membrane | PF02630.9(SCO1-SenC) | 2,725 | 3,735 |
|  | Bcin16g04050 | nd | NADP-specific glutamate dehydrogenase | PF02812.13(ELFV_dehydrog_N) + PF00208.16(ELFV_dehydrog) | 2,750 | 4,263 |
|  | Bcin02g01770 | nd | NAD NADP octopine nopaline dehydrogenase | PF01210.18(NAD_Gly3P_dh_N) + PF02317.12(Octopine_DH) | 2,811 | 3,029 |
|  | Bcin14g00210 | nd | nadh oxidase | PF00724.15(Oxidored_FMN) | 2,815 | 15,203 |
|  | Bcin14g00100 | nd | NADH:ubiquinone B18 subunit | PF05676.8(NDUF_B7) | 2,878 | 1,778 |
|  | Bcin05g03840 | nd | nd | nd | 2,887 | 1,618 |
|  | Bcin01g03710 | nd | Acyl- N-acyltransferase | PF13523.1(Acetyltransf_8) | 2,895 | 1,885 |
|  | Bcin03g07670 | nd | nad-specific glutamate dehydrogenase | PF05088.7(Bac_GDH) + PF00208.16(ELFV_dehydrog) | 3,057 | 1,865 |
|  | Bcin05g02460 | nd | bax inhibitor family | PF01027.15(Bax1-I) | 3,091 | 4,345 |
|  | Bcin03g03910 | nd | sucrase ferredoxin domain containing | PF06999.7(Suc_Fer-like) | 3,124 | 9,071 |
|  | Bcin11g00160 | nd | adenylyl-sulfate kinase | PF01583.15(APS_kinase) | 3,154 | 2,264 |
|  | Bcin12g03650 | nd | norsolorinic acid reductase | PF00248.16(Aldo_ket_red) | 3,685 | 9,249 |
|  | Bcin14g00850 | BcPga1 | polygalacturonase partial | PF00295.12(Glyco_hydro_28) | 3,721 | 2,534 |
|  | Bcin01g06010 | nd | glycoside hydrolase family 16 | PF00722.16(Glyco_hydro_16) | 3,803 | 3,635 |
|  | Bcin06g07480 | nd | tam domain methyltransferase | PF13489.1(Methyltransf_23) | 3,837 | 8,662 |
|  | Bcin06g06040 | nd | nd | PF03856.8(SUN) | 3,890 | 1,990 |
|  | Bcin12g04980 | BcNrps1 | nonribosomal peptide synthase -like | PF00550.20(PP-binding) + PF00668.15(Condensation) + PF00501.23(AMP-binding) + PF00550.20(PP-binding) + PF00668.15(Condensation) | 4,763 | 17,918 |
|  | Bcin01g03270 | nd | nd | PF05368.8(NmrA) | 5,043 | 5,389 |
|  | Bcin05g03230 | nd | phosphoketolase | PF09364.5(XFP_N) + PF03894.10(XFP) + PF09363.5(XFP_C) | 5,092 | 4,848 |
|  | Bcin07g00300 | nd | nd | PF01814.18(Hemerythrin) | 5,505 | 9,864 |
|  | Bcin05g07140 | nd | mitochondrial import inner membrane translocase subunit TIM10 | PF02953.10(zf-Tim10_DDP) | 5,916 | 2,184 |
|  | Bcin13g03410 | nd | paf acetylhydrolase family | PF12695.2(Abhydrolase_5) | 6,920 | 26,291 |
|  | Bcin14g02940 | nd | nd | PF00248.16(Aldo_ket_red) | 8,539 | 21,077 |
|  | Bcin12g04900 | nd | Salicylate hydroxylase | PF01494.14(FAD_binding_3) | 11,896 | 23,193 |
|  | Bcin10g02140* | nd | ABC transporter | PF00664.18(ABC_membrane) + PF00005.22(ABC_tran) + PF00664.18(ABC_membrane) + PF00005.22(ABC_tran) | 1,580 | 2,880 |
|  | Bcin16g00002* | nd | FAD-dependent pyridine nucleotide-disulfide partial | PF13738.1(Pyr_redox_3) | 1,600 | 5,800 |
|  | Bcin09g05930* | nd | ABC transporter | PF00664.18(ABC_membrane) + PF00005.22(ABC_tran) + PF00664.18(ABC_membrane) + PF00005.22(ABC_tran) | 1,900 | 3,600 |
|  | Bcin11g01080* | nd | nd | PF00078.22(RVT_1) | 2,364 | 3,091 |
|  | Bcin14g03160* | BcGst5 | Glutathione S-transferase | PF13417.1(GST_N_3) + PF00043.20(GST_C) | 3,000 | 15,000 |
|  | Bcin01g11540* | nd | aldolase | PF03328.9(HpcH_HpaI) | 4,500 | 14,500 |
|  | Bcin02g01990* | nd | nd | PF13460.1(NAD_binding_10) | 6,067 | 5,600 |
|  | Bcin12g04940* | nd | cytochrome P450 monooxygenase | PF00067.17(p450) | 6,500 | 8,667 |
|  | Bcin06g03440* | BcAox | alternative oxidase | PF01786.12(AOX) | 6,750 | 6,750 |
|  | Bcin09g06120* | nd | zinc-regulated transporter 1 | PF02535.17(Zip) | 7,500 | 14,000 |
|  | Bcin13g00710* | BcAtrB | abc transporter | PF14510.1(ABC_trans_N) + PF00005.22(ABC_tran) + PF01061.19(ABC2_membrane) + PF06422.7(PDR_CDR) + PF00005.22(ABC_tran) + PF01061.19(ABC2_membrane) | 9,000 | 43,000 |
|  | Bcin14g05580* | nd | 2-polyprenyl-6-methoxyphenol hydroxylase | PF01494.14(FAD_binding_3) | 19,333 | 17,667 |

| **Bos1+ / Sak1+** | **Accession number** | **Name** | **Description** | **Pfam** | ***Δbos1* / WT** | ***Δsak1* / WT** |
| --- | --- | --- | --- | --- | --- | --- |
|  | Bcin07g05550 | nd | nd | nd | 0,065 | 0,033 |
|  | Bcin14g04090 | nd | nd | PF02036.12(SCP2) | 0,066 | 0,058 |
|  | Bcin01g08160 | nd | Pesticidal crystal cry6Aa | nd | 0,085 | 0,091 |
|  | Bcin06g02510 | nd | alpha beta-hydrolase | PF00450.17(Peptidase_S10) | 0,117 | 0,108 |
|  | Bcin12g06430 | nd | retinol dehydrogenase 8 | PF00106.20(adh_short) | 0,117 | 0,393 |
|  | Bcin03g01920 | BcCat5 | Catalase | PF00199.14(Catalase) + PF06628.7(Catalase-rel) | 0,120 | 0,039 |
|  | Bcin15g03150 | nd | tripeptidyl-peptidase 1 precursor | PF09286.6(Pro-kuma_activ) + PF00082.17(Peptidase_S8) | 0,126 | 0,040 |
|  | Bcin16g03540 | nd | acetolactate synthase | PF02776.13(TPP_enzyme_N) + PF00205.17(TPP_enzyme_M) + PF02775.16(TPP_enzyme_C) | 0,136 | 0,210 |
|  | Bcin07g05560 | nd | nd | nd | 0,145 | 0,080 |
|  | Bcin01g02910 | nd | histidine acid phosphatase | PF00328.17(His_Phos_2) | 0,158 | 0,169 |
|  | Bcin05g01520 | nd | glycoside hydrolase family 16 | PF00722.16(Glyco_hydro_16) | 0,159 | 0,105 |
|  | Bcin14g04580 | nd | chlorophyll synthesis pathway | PF08240.7(ADH_N) + PF00107.21(ADH_zinc_N) | 0,160 | 0,009 |
|  | Bcin02g09370 | nd | C2 domain-containing | PF00168.25(C2) + PF00168.25(C2) | 0,165 | 0,130 |
|  | Bcin04g04190 | nd | glycoside hydrolase family 15 | PF00723.16(Glyco_hydro_15) + PF00686.14(CBM_20) | 0,169 | 0,126 |
|  | Bcin10g00030 | nd | dimeric alpha-beta barrel | PF07110.6(EthD) | 0,182 | 0,012 |
|  | Bcin01g08150 | nd | nd | nd | 0,185 | 0,169 |
|  | Bcin04g06450 | nd | aldo-keto reductase (AKR13) | PF00248.16(Aldo_ket_red) | 0,188 | 0,199 |
|  | Bcin13g04640 | nd | short chain dehydrogenase reductase | PF00106.20(adh_short) | 0,202 | 0,029 |
|  | Bcin01g02880 | nd | alcohol dehydrogenase ii | PF08240.7(ADH_N) + PF00107.21(ADH_zinc_N) | 0,203 | 0,144 |
|  | Bcin12g03390 | nd | glycoside hydrolase family 15 | PF00723.16(Glyco_hydro_15) + PF00686.14(CBM_20) | 0,204 | 0,053 |
|  | Bcin03g01040 | nd | glutamate decarboxylase | PF00282.14(Pyridoxal_deC) | 0,206 | 0,331 |
|  | Bcin02g03090 | nd | glycoside hydrolase family 1 | PF00232.13(Glyco_hydro_1) | 0,222 | 0,084 |
|  | Bcin16g04640 | nd | formate dehydrogenase | PF00389.25(2-Hacid_dh) + PF02826.14(2-Hacid_dh_C) | 0,236 | 0,169 |
|  | Bcin11g02720 | nd | alcohol dehydrogenase | PF00248.16(Aldo_ket_red) | 0,237 | 0,040 |
|  | Bcin16g02700 | nd | dihydroxy-acid dehydratase | PF00920.16(ILVD_EDD) + PF00920.16(ILVD_EDD) | 0,243 | 0,599 |
|  | Bcin15g03620 | nd | glycogen phosphorylase | PF00343.15(Phosphorylase) | 0,256 | 0,257 |
|  | Bcin12g01020 | BcOah | oxaloacetate partial | PF13714.1(PEP_mutase) | 0,263 | 0,010 |
|  | Bcin02g01610 | BcNde3 | apoptosis-inducing factor 1 | PF07992.9(Pyr_redox_2) + PF00070.22(Pyr_redox) | 0,264 | 0,110 |
|  | Bcin10g01030 | BcPrx9 | TSA family | PF08534.5(Redoxin) | 0,265 | 0,055 |
|  | Bcin15g03640 | nd | 6-hydroxy-d-nicotine oxidase | PF01565.18(FAD_binding_4) + PF08031.7(BBE) | 0,274 | 0,113 |
|  | Bcin11g00140 | nd | nd | nd | 0,276 | 0,262 |
|  | Bcin04g05680 | nd | aldehyde dehydrogenase | PF00171.17(Aldedh) | 0,282 | 0,568 |
|  | Bcin01g10310 | nd | glycogen debranching enzyme | PF14699.1(hGDE_N) + PF14701.1(hDGE_amylase) + PF14702.1(hGDE_central) + PF06202.9(GDE_C) | 0,283 | 0,234 |
|  | Bcin11g06440 | nd | glycoside hydrolase family 31 | PF01055.21(Glyco_hydro_31) | 0,285 | 0,201 |
|  | Bcin16g02810 | nd | nd | nd | 0,299 | 0,510 |
|  | Bcin06g04940 | nd | short-chain dehydrogenase | PF00106.20(adh_short) | 0,307 | 0,099 |
|  | Bcin08g02430 | nd | uracil phosphoribosyltransferase | PF14681.1(UPRTase) | 0,313 | 0,227 |
|  | Bcin06g00330 | nd | tripeptidyl-peptidase 1 precursor | PF09286.6(Pro-kuma_activ) + PF00082.17(Peptidase_S8) | 0,313 | 0,152 |
|  | Bcin08g03150 | nd | dihydroxyacetone kinase | PF02733.12(Dak1) + PF02734.12(Dak2) | 0,324 | 0,239 |
|  | Bcin09g07140 | nd | hypothetical protein BcDW1_7862 | nd | 0,327 | 0,634 |
|  | Bcin02g00013 | nd | salicylate 1-monooxygenase | PF01494.14(FAD_binding_3) | 0,336 | 0,158 |
|  | Bcin06g02760 | nd | nd | PF07978.8(NIPSNAP) + PF07978.8(NIPSNAP) | 0,356 | 0,557 |
|  | Bcin06g06350 | BcNqo1 | minor allergen alt a 7 | PF00258.20(Flavodoxin_1) | 0,362 | 0,410 |
|  | Bcin04g00600 | nd | P-loop containing nucleoside triphosphate hydrolase | PF00485.13(PRK) | 0,362 | 0,261 |
|  | Bcin01g03390 | nd | glycosyl hydrolase family 38 | PF01074.17(Glyco_hydro_38) + PF09261.6(Alpha-mann_mid) + PF07748.8(Glyco_hydro_38C) | 0,364 | 0,180 |
|  | Bcin10g02340 | nd | s-formylglutathione hydrolase | PF00756.15(Esterase) | 0,373 | 0,548 |
|  | Bcin11g04130 | nd | cyanamide hydratase | PF01966.17(HD) | 0,374 | 0,625 |
|  | Bcin09g00650 | nd | 2-methylcitrate synthase | PF00285.16(Citrate_synt) | 0,380 | 0,399 |
|  | Bcin15g04670 | BcSer8 | nd | PF09286.6(Pro-kuma_activ) + PF00082.17(Peptidase_S8) | 0,381 | 0,132 |
|  | Bcin14g00480 | nd | glycoside hydrolase family 3 | PF00933.16(Glyco_hydro_3) + PF01915.17(Glyco_hydro_3_C) + PF07691.7(PA14) + PF14310.1(Fn3-like) | 0,383 | 0,225 |
|  | Bcin09g02680 | nd | glycoside hydrolase family 63 | PF03200.11(Glyco_hydro_63) | 0,387 | 0,254 |
|  | Bcin15g02090 | nd | 1,4-alpha-glucan-branching enzyme | PF02922.13(CBM_48) + PF00128.19(Alpha-amylase) + PF02806.13(Alpha-amylase_C) | 0,394 | 0,239 |
|  | Bcin07g03170 | nd | cell lysis | PF02453.12(Reticulon) | 0,396 | 0,547 |
|  | Bcin03g06200 | nd | GTP cyclohydrolase II | PF12471.3(GTP_CH_N) + PF00925.15(GTP_cyclohydro2) | 0,398 | 0,457 |
|  | Bcin08g02390 | nd | Peptidase S28 | PF05577.7(Peptidase_S28) | 0,401 | 0,151 |
|  | Bcin07g04370 | nd | peptidase s41 family | nd | 0,406 | 0,099 |
|  | Bcin02g05580 | nd | nd | nd | 0,407 | 0,472 |
|  | Bcin01g10150 | BcPsd | phosphatidylserine decarboxylase | PF02666.10(PS_Dcarbxylase) | 0,418 | 0,320 |
|  | Bcin09g07130 | nd | nd | nd | 0,421 | 0,564 |
|  | Bcin02g01070 | nd | glucosidase ii alpha subunit | PF13802.1(Gal_mutarotas_2) + PF01055.21(Glyco_hydro_31) | 0,421 | 0,502 |
|  | Bcin03g03440 | nd | haloacid dehalogenase-like hydrolase | PF13419.1(HAD_2) | 0,423 | 0,088 |
|  | Bcin08g02080 | nd | antibiotic biosynthesis monooxygenase | PF03992.11(ABM) | 0,426 | 0,375 |
|  | Bcin07g02370 | nd | Class I glutamine amidotransferase | PF13278.1(DUF4066) | 0,428 | 0,528 |
|  | Bcin11g03260 | BcPio9 | nd | PF11720.3(Inhibitor_I78) | 0,429 | 0,393 |
|  | Bcin11g04710 | nd | glycerol dehydrogenase | PF00248.16(Aldo_ket_red) | 0,440 | 0,562 |
|  | Bcin06g01920 | nd | annexin anxc4 | PF00191.15(Annexin) | 0,442 | 0,388 |
|  | Bcin16g02220 | nd | lysine decarboxylase | PF03641.9(Lysine_decarbox) | 0,443 | 0,394 |
|  | Bcin02g02910 | nd | Pentulose kinase | PF00370.16(FGGY_N) + PF02782.11(FGGY_C) | 0,455 | 0,298 |
|  | Bcin02g04670 | nd | pyroglutamyl peptidase type | PF01470.12(Peptidase_C15) | 0,461 | 0,511 |
|  | Bcin01g10100 | BcPic7 | pyruvate kinase | PF00224.16(PK) + PF02887.11(PK_C) | 0,465 | 0,348 |
|  | Bcin04g05300 | BcGlr1 | glutathione-disulfide reductase | PF07992.9(Pyr_redox_2) + PF00070.22(Pyr_redox) + PF02852.17(Pyr_redox_dim) | 0,473 | 0,360 |
|  | Bcin01g00750 | nd | DUF718 domain-containing | PF05336.8(DUF718) | 0,492 | 0,296 |
|  | Bcin12g04970 | nd | FAD NAD(P)-binding domain-containing | PF01494.14(FAD_binding_3) | 0,494 | 0,548 |
|  | Bcin03g02930 | BcCla4 | mitogen-activated kinase : p21-activated kinase (PAK) | PF00786.23(PBD) + PF00069.20(Pkinase) | 0,495 | 0,640 |
|  | Bcin11g00830 | nd | FAD NAD(P)-binding domain-containing | PF13738.1(Pyr_redox_3) | 0,501 | 0,468 |
|  | Bcin02g05270 | nd | udp-glucose:glyco glucosyltransferase | PF06427.6(UDP-g_GGTase) | 0,503 | 0,500 |
|  | Bcin16g01550 | nd | nd | nd | 0,506 | 0,454 |
|  | Bcin06g03060 | BcGst9 | glutathione s- | PF02798.15(GST_N) + PF14497.1(GST_C_3) | 0,520 | 0,232 |
|  | Bcin05g01260 | nd | nd | nd | 0,523 | 0,462 |
|  | Bcin09g02460 | nd | cytochrome b5 | PF00173.23(Cyt-b5) | 0,523 | 0,449 |
|  | Bcin01g11330 | nd | glucosidase 2 subunit beta | PF12999.2(PRKCSH-like) + PF12999.2(PRKCSH-like) + PF13015.1(PRKCSH_1) | 0,527 | 0,590 |
|  | Bcin03g02420 | nd | nd | nd | 0,530 | 0,410 |
|  | Bcin11g05710 | nd | oligosaccharyltransferase alpha subunit | PF04597.9(Ribophorin_I) | 0,532 | 0,640 |
|  | Bcin15g04410 | nd | metallopeptidase | PF01432.15(Peptidase_M3) | 0,534 | 0,531 |
|  | Bcin04g05230 | nd | mitochondrial chaperone Frataxin | PF01491.11(Frataxin_Cyay) | 0,534 | 0,468 |
|  | Bcin01g01260 | nd | DJ-1 -type | PF13278.1(DUF4066) | 0,536 | 0,399 |
|  | Bcin09g02790 | nd | acetate-- ligase | PF00501.23(AMP-binding) + PF13193.1(AMP-binding_C) | 0,538 | 0,456 |
|  | Bcin04g06850 | nd | retinol dehydrogenase | PF00106.20(adh_short) | 0,538 | 0,472 |
|  | Bcin09g02470 | nd | hypothetical protein BC1G_07135 | nd | 0,543 | 0,629 |
|  | Bcin12g01870 | nd | quinone oxidoreductase | PF08240.7(ADH_N) + PF00107.21(ADH_zinc_N) | 0,553 | 0,350 |
|  | Bcin03g06220 | nd | uracil phosphoribosyltransferase | PF14681.1(UPRTase) | 0,558 | 0,427 |
|  | Bcin12g02430 | nd | Pyridoxal phosphate-dependent major subdomain 2 | PF13500.1(AAA_26) + PF00202.16(Aminotran_3) + PF00202.16(Aminotran_3) | 0,562 | 0,235 |
|  | Bcin02g03490 | nd | cystathionine gamma-lyase | PF01053.15(Cys_Met_Meta_PP) | 0,576 | 0,316 |
|  | Bcin06g04660 | BcPio12, BcGar1 | aldo keto reductase | PF00248.16(Aldo_ket_red) | 0,579 | 0,644 |
|  | Bcin08g02380 | nd | disulfide-isomerase erp38 | PF00085.15(Thioredoxin) + PF00085.15(Thioredoxin) + PF07749.7(ERp29) | 0,580 | 0,640 |
|  | Bcin11g05700 | BcHxk | hexokinase | PF00349.16(Hexokinase_1) + PF03727.11(Hexokinase_2) | 0,581 | 0,484 |
|  | Bcin02g06170 | nd | nuclear condensin complex subunit smc4 | PF02463.14(SMC_N) + PF06470.8(SMC_hinge) | 0,583 | 0,417 |
|  | Bcin04g06400 | nd | glycerol-3-phosphate dehydrogenase | PF01266.19(DAO) | 0,589 | 0,557 |
|  | Bcin05g05290 | nd | nd | PF05368.8(NmrA) | 0,592 | 0,032 |
|  | Bcin08g05100 | nd | 3-methylcrotonyl- carboxylase subunit alpha | PF00289.17(CPSase_L_chain) + PF02786.12(CPSase_L_D2) + PF02785.14(Biotin_carb_C) + PF00364.17(Biotin_lipoyl) | 0,595 | 0,364 |
|  | Bcin08g05630 | nd | cyanate hydratase | PF02560.9(Cyanate_lyase) | 0,600 | 0,553 |
|  | Bcin13g05580 | nd | alcohol dehydrogenase | PF08240.7(ADH_N) + PF00107.21(ADH_zinc_N) | 0,608 | 0,418 |
|  | Bcin14g04770 | nd | PSP1 domain-containing | PF04468.7(PSP1) | 0,609 | 0,512 |
|  | Bcin16g03410 | nd | actin depolymerizing | PF00241.15(Cofilin_ADF) + PF00241.15(Cofilin_ADF) | 0,609 | 0,472 |
|  | Bcin16g04800 | nd | malate dehydrogenase | PF00056.18(Ldh_1_N) + PF02866.13(Ldh_1_C) | 0,617 | 0,601 |
|  | Bcin14g01030 | nd | S-(hydroxymethyl)glutathione dehydrogenase | PF08240.7(ADH_N) + PF00107.21(ADH_zinc_N) | 0,617 | 0,548 |
|  | Bcin09g01930 | nd | 2-methylcitrate dehydratase | PF03972.9(MmgE_PrpD) | 0,620 | 0,601 |
|  | Bcin03g08100 | BcBrn2 | tetrahydroxynaphthalene reductase | PF00106.20(adh_short) | 0,622 | 0,007 |
|  | Bcin15g04970 | nd | glucose-6-phosphate isomerase | PF00342.14(PGI) | 0,629 | 0,588 |
|  | Bcin02g07690 | nd | enoyl- hydratase isomerase | PF00378.15(ECH) | 0,631 | 0,583 |
|  | Bcin03g08240 | nd | acyl binding | PF00887.14(ACBP) | 0,635 | 0,608 |
|  | Bcin08g06300 | nd | 6-phosphofructokinase | PF00365.15(PFK) + PF00365.15(PFK) | 0,635 | 0,591 |
|  | Bcin03g05490 | nd | hydroxyacylglutathione hydrolase | PF00753.22(Lactamase_B) | 0,637 | 0,655 |
|  | Bcin09g00510 | nd | Chaperone J-domain-containing | PF00226.26(DnaJ) + PF00684.14(DnaJ_CXXCXGXG) + PF01556.13(CTDII) | 0,638 | 0,556 |
|  | Bcin02g06130 | nd | nd | PF01237.13(Oxysterol_BP) + PF01237.13(Oxysterol_BP) | 0,644 | 0,489 |
|  | Bcin04g00570 | BcPrx1 | merozoite capping -1 | PF00578.16(AhpC-TSA) | 0,647 | 0,617 |
|  | Bcin03g01520 | nd | NADP-dependent alcohol dehydrogenase | PF08240.7(ADH_N) + PF00107.21(ADH_zinc_N) | 0,653 | 0,017 |
|  | Bcin03g01560 | nd | aldo keto reductase | PF00248.16(Aldo_ket_red) | 0,654 | 0,288 |
|  | Bcin16g01950 | nd | glycoside hydrolase family 63 | PF03200.11(Glyco_hydro_63) | 0,655 | 0,601 |
|  | Bcin02g04770 | nd | methionine vitamin-b12 | PF01717.13(Meth_synt_2) | 0,658 | 0,179 |
|  | Bcin03g00010* | nd | nd | nd | 0,000 | 0,000 |
|  | Bcin11g02630* | nd | phytanoyl- dioxygenase | PF05721.8(PhyH) | 0,050 | 0,025 |
|  | Bcin04g05000* | BcPpoB-like, PpoA90 | fatty acid oxygenase | PF03098.10(An_peroxidase) + PF00067.17(p450) | 0,085 | 0,122 |
|  | Bcin03g00005* | nd | Aegerolysin family | PF06355.8(Aegerolysin) | 0,118 | 0,265 |
|  | Bcin02g05500* | nd | alpha beta-hydrolase | PF12697.2(Abhydrolase_6) | 0,154 | 0,115 |
|  | Bcin01g10130* | nd | cytosolic nonspecific dipeptidase | PF01546.23(Peptidase_M20) + PF07687.9(M20_dimer) | 0,161 | 0,161 |
|  | Bcin07g05540* | nd | nd | nd | 0,169 | 0,152 |
|  | Bcin05g07640* | nd | cytochrome P450 | PF00067.17(p450) | 0,188 | 0,250 |
|  | Bcin15g03060* | nd | nd | PF06101.6(DUF946) | 0,215 | 0,046 |
|  | Bcin11g02670* | nd | gluconate 5-dehydrogenase | PF00106.20(adh_short) | 0,233 | 0,100 |
|  | Bcin11g02700* | BcPks4 | polyketide synthase | PF00109.21(ketoacyl-synt) + PF02801.17(Ketoacyl-synt_C) + PF00698.16(Acyl_transf_1) + PF14765.1(PS-DH) + PF08242.7(Methyltransf_12) + PF08659.5(KR) + PF00550.20(PP-binding) | 0,258 | 0,169 |
|  | Bcin12g06180* | nd | cyanide hydratase | PF00795.17(CN_hydrolase) | 0,261 | 0,435 |
|  | Bcin05g01530* | nd | glycosyltransferase family 2 | PF13632.1(Glyco_trans_2_3) | 0,269 | 0,654 |
|  | Bcin12g00300* | nd | Glycoside family 35 | PF01301.14(Glyco_hydro_35) + PF10435.4(BetaGal_dom2) + PF13363.1(BetaGal_dom3) + PF13364.1(BetaGal_dom4_5) + PF13364.1(BetaGal_dom4_5) | 0,276 | 0,276 |
|  | Bcin13g01770* | nd | nd | nd | 0,276 | 0,172 |
|  | Bcin08g00280* | nd | carboxypeptidase s1 | PF00450.17(Peptidase_S10) | 0,281 | 0,125 |
|  | Bcin01g08340* | nd | (sam-dependent methyltransferases) | PF08242.7(Methyltransf_12) | 0,300 | 0,150 |
|  | Bcin14g00090* | nd | thermophilic desulfurizing enzyme family | PF02771.11(Acyl-CoA_dh_N) + PF08028.6(Acyl-CoA_dh_2) | 0,318 | 0,273 |
|  | Bcin10g00040* | BcPks7 | hybrid pks-nrps | PF00109.21(ketoacyl-synt) + PF02801.17(Ketoacyl-synt_C) + PF00698.16(Acyl_transf_1) + PF14765.1(PS-DH) + PF08242.7(Methyltransf_12) + PF08659.5(KR) + PF00550.20(PP-binding) + PF00668.15(Condensation) + PF00501.23(AMP-binding) + PF00550.20(PP-binding) + PF07993.7(NAD_binding_4) | 0,321 | 0,113 |
|  | Bcin03g03480* | nd | glycoside hydrolase family 10 | PF00734.13(CBM_1) + PF00331.15(Glyco_hydro_10) | 0,400 | 0,200 |
|  | Bcin14g00650* | nd | glycoside hydrolase family 31 | PF13802.1(Gal_mutarotas_2) + PF01055.21(Glyco_hydro_31) | 0,404 | 0,213 |
|  | Bcin03g05040* | nd | rna interference and gene silencing | PF08699.5(DUF1785) + PF02170.17(PAZ) + PF02171.12(Piwi) | 0,476 | 0,622 |
|  | Bcin11g02730* | BcPio5 | CDP-alcohol phosphatidyltransferase | PF13344.1(Hydrolase_6) + PF13242.1(Hydrolase_like) | 0,619 | 0,155 |

| **Bos1- / Sak1+** | **Accession number** | **Name** | **Description** | **Pfam** | ***Δbos1* / WT** | ***Δsak1* / WT** |
| --- | --- | --- | --- | --- | --- | --- |
|  | Bcin06g05640 | nd | RNA polymerase Rpb3 insert domain-containing | PF01193.19(RNA_pol_L) + PF01000.21(RNA_pol_A_bac) | 1,518 | 0,560 |
|  | Bcin12g00190 | nd | GMP synthase | PF00117.23(GATase) + PF02540.12(NAD_synthase) + PF00958.17(GMP_synt_C) | 1,546 | 0,607 |
|  | Bcin05g04910 | nd | immunogenic | PF15406.1(PH_6) | 1,559 | 0,656 |
|  | Bcin01g06060 | nd | nd | nd | 1,983 | 0,118 |
|  | Bcin03g01540 | nd | GMC oxidoreductase | PF00732.14(GMC_oxred_N) + PF05199.8(GMC_oxred_C) | 1,996 | 0,192 |
|  | Bcin15g00100 | nd | nd | nd | 2,040 | 0,130 |
|  | Bcin01g11220 | nd | glycoside hydrolase family 17 | PF00332.13(Glyco_hydro_17) | 2,191 | 0,519 |
|  | Bcin13g01570 | nd | zinc finger zpr1 | PF03367.8(zf-ZPR1) + PF03367.8(zf-ZPR1) | 2,739 | 0,613 |
|  | Bcin04g05700 | nd | alcohol dehydrogenase | PF08240.7(ADH_N) + PF00107.21(ADH_zinc_N) | 3,067 | 0,114 |
|  | Bcin01g00010 | BcBoA1 | nd | PF05368.8(NmrA) | 3,381 | 0,525 |
|  | Bcin14g02510 | BcLcc2 | laccase 2 | PF07732.10(Cu-oxidase_3) + PF00394.17(Cu-oxidase) + PF07731.9(Cu-oxidase_2) | 3,867 | 0,294 |
|  | Bcin14g03430* | nd | pectin lyase a precursor | PF00544.14(Pec_lyase_C) | 1,600 | 0,200 |
|  | Bcin16g00930* | CND3 | nd | nd | 1,900 | 0,550 |
|  | Bcin14g01290* | BcPks11 | polyketide synthase | PF00109.21(ketoacyl-synt) + PF02801.17(Ketoacyl-synt_C) + PF00698.16(Acyl_transf_1) + PF14765.1(PS-DH) + PF08242.7(Methyltransf_12) + PF00107.21(ADH_zinc_N) + PF08659.5(KR) + PF00550.20(PP-binding) | 3,417 | 0,333 |

| **Bos1+ / Sak1-** | **Accession number** | **Name** | **Description** | **Pfam** | ***Δbos1* / WT** | ***Δsak1* / WT** |
| --- | --- | --- | --- | --- | --- | --- |
|  | Bcin12g06370 | BcBot4 | benzoate 4-monooxygenase cytochrome p450 | PF00067.17(p450) | 0,055 | 1,536 |
|  | Bcin12g06400 | BcBot3 | cytochrome P450 monooxygenase | PF00067.17(p450) | 0,134 | 1,829 |
|  | Bcin01g10380 | nd | NADPH--cytochrome P450 reductase | PF00067.17(p450) + PF00258.20(Flavodoxin_1) + PF00667.15(FAD_binding_1) + PF00175.16(NAD_binding_1) | 0,284 | 32,005 |
|  | Bcin06g06240 | nd | taurine catabolism dioxygenase | PF02668.11(TauD) | 0,418 | 3,736 |
|  | Bcin12g06390 | BcBot2 | Presilphiperfolan-8-beta-ol synthase | PF03936.11(Terpene_synth_C) | 0,514 | 1,895 |
|  | Bcin01g11450 | BcNrps7 | nonribosomal siderophore peptide synthase | PF00501.23(AMP-binding) + PF13193.1(AMP-binding_C) + PF00668.15(Condensation) | 0,535 | 10,278 |
|  | Bcin05g04870 | nd | flavocytochrome c | PF00890.19(FAD_binding_2) + PF00173.23(Cyt-b5) | 0,537 | 1,586 |
|  | Bcin02g07220 | nd | acyl- dehydrogenase | PF00173.23(Cyt-b5) + PF02771.11(Acyl-CoA_dh_N) + PF02770.14(Acyl-CoA_dh_M) + PF00441.19(Acyl-CoA_dh_1) | 0,595 | 6,826 |
|  | Bcin13g00770 | nd | epoxide hydrolase | PF12697.2(Abhydrolase_6) | 0,626 | 2,354 |
|  | Bcin12g03480 | nd | phosphatidylethanolamine methyltransferase | PF04191.8(PEMT) + PF04191.8(PEMT) | 0,631 | 2,206 |
|  | Bcin09g03440 | nd | integral membrane DUF92 | PF01940.11(DUF92) | 0,645 | 2,719 |
|  | Bcin12g06460 | nd | dienelactone hydrolase | PF01738.13(DLH) | 0,659 | 1,886 |
|  | Bcin04g03720* | nd | salicylate hydroxylase | PF01494.14(FAD_binding_3) + PF01494.14(FAD_binding_3) | 0,583 | 4,167 |
